# Release of P-TEFb from the Super Elongation Complex promotes HIV-1 latency reversal

**DOI:** 10.1101/2024.03.01.582881

**Authors:** William J. Cisneros, Miriam Walter, Shimaa H.A. Soliman, Lacy M. Simons, Daphne Cornish, Ariel W. Halle, Eun-Young Kim, Steven M. Wolinsky, Ali Shilatifard, Judd F. Hultquist

## Abstract

The persistence of HIV-1 in long-lived latent reservoirs during suppressive antiretroviral therapy (ART) remains one of the principal barriers to a functional cure. Blocks to transcriptional elongation play a central role in maintaining the latent state, and several latency reversal strategies focus on the release of positive transcription elongation factor b (P-TEFb) from sequestration by negative regulatory complexes, such as the 7SK complex and BRD4. Another major cellular reservoir of P-TEFb is in Super Elongation Complexes (SECs), which play broad regulatory roles in host gene expression. Still, it is unknown if the release of P-TEFb from SECs is a viable latency reversal strategy. Here, we demonstrate that the SEC is not required for HIV-1 replication in primary CD4+ T cells and that a small molecular inhibitor of the P-TEFb/SEC interaction (termed KL-2) increases viral transcription. KL-2 acts synergistically with other latency reversing agents (LRAs) to reactivate viral transcription in several cell line models of latency in a manner that is, at least in part, dependent on the viral Tat protein. Finally, we demonstrate that KL-2 enhances viral reactivation in peripheral blood mononuclear cells (PBMCs) from people living with HIV on suppressive ART, most notably in combination with inhibitor of apoptosis protein antagonists (IAPi). Taken together, these results suggest that the release of P-TEFb from cellular SECs may be a novel route for HIV-1 latency reactivation.

**AUTHOR SUMMARY:** Since the start of the HIV pandemic, it is estimated that nearly 86 million people have been infected with the virus, and about 40 million people have died. Modern antiretroviral therapies potently restrict viral replication and prevent the onset of AIDS, saving millions of lives. However, these therapies are not curative due to the persistence of the virus in a silenced or ‘latent’ state in long-lived cells of the body. One proposed strategy to clear this latent reservoir, termed “shock and kill”, is to activate these silenced viruses such that the infected cells can be cleared from the body by the immune system. While several drugs have been developed that can activate latent viruses, none have proven effective at reducing the size of the latent reservoir in patients in clinical trials. Here, we describe a new method for latency reactivation using a small molecule inhibitor of a human protein complex called the Super Elongation Complex (SEC). Inhibiting the SEC enhances viral transcription during active infection and triggers the reactivation of latent viruses, especially when in combination with other latency reversing agents. These results pave the way for developing more effective strategies to reactivate latent viruses towards a functional cure.

## INTRODUCTION

The persistence of replication-competent, but transcriptionally inhibited HIV-1 proviral DNA in long-lived latent cellular reservoirs is a significant barrier to the development of a functional cure (1, 2) Even after long-term suppressive antiretroviral therapy (ART), spontaneous reactivation of proviral gene expression from the latent reservoir is sufficient to initiate viral rebound shortly after ART cessation, thus requiring life-long adherence (3–5). Multifaceted and heterogenous blocks to viral gene expression establish and maintain HIV-1 latency at the epigenetic, transcriptional, and post-transcriptional levels (2). Several strategies to either reactivate latent proviruses and clear infected cells (*i.e.*, “shock and kill”) or reinforce latency to prevent spontaneous reactivation (*i.e.*, “block and lock”) are currently under investigation (6, 7).

While many small molecule latency reversing agents (LRAs) have been described that reactivate latent proviruses *ex vivo*, they have had little success in clinical trials (8, 9). This failure is partly due to the latent reservoir’s heterogenous nature, such that treatment by a single agent that acts to target a specific block may only ever reactivate a small fraction of proviruses *in vivo* (10). Even if transcriptional reactivation is achieved, it is unlikely this is sufficient to result in the clearance of these infected cells without additional immune augmentation such as the administration of antibodies, vaccines, or immunotherapies to prime the immune system for appropriate and potent responses (6). Given these limitations, much research is now focused on combinatorial approaches to trigger more widespread and robust reactivation (11). For example, a recent study reported synergistic reactivation potential between an activator of non-canonical NFkB signaling (AZD5582) and BET bromodomain inhibitors that act to lift blocks to transcriptional initiation and elongation, respectively (12).

Blocks to transcriptional elongation are major contributors to establishing and maintaining HIV-1 latency (13, 14). After integration of the proviral DNA, RNA Polymerase II (RNA Pol II) is recruited to the transcription start site by transcription factors that bind *cis-*elements in the HIV-1 promoter region. After transcriptional initiation, RNA Pol II synthesizes 20-60 nucleotides before stalling through a well-conserved process known as promoter-proximal pausing (15). Pausing is enforced by several negative elongation factors, including negative elongation factor (NELF), DRB Sensitivity Inducing Factor (DSIF), and the RNA Polymerase II Associated Factor 1 (PAF1) complex (16–18). Pause release is regulated by positive transcription elongation factor-b (P-TEFb), a heterodimeric protein complex composed of cyclin-dependent kinase 9 (CDK9) and cyclin T1 (CCNT1) (19–21). P-TEFb phosphorylates the C-terminal tail of RNA Pol II and several negative elongation factors, which collectively license transcriptional elongation (22–24).

Recruitment of P-TEFb to sites of nascent transcription is a highly regulated process mediated by several cellular complexes. The majority of cellular P-TEFb is sequestered in an inactive state by the 7SK ribonucleoprotein (RNP) complex (25, 26). Diverse extracellular stimuli and intracellular signals can induce the release of P-TEFb from the 7SK (27–29) where it can be recruited to sites of nascent transcription by transcription factors (*i.e.*, NFkB and c-MYC) (30–32), epigenetic regulators (*i.e.*, BRD4) (33, 34), or super elongation complexes (SECs) composed of an ARF4/FMR2 (AFF) family scaffold protein in complex with AF9, ENL, an eleven-nineteen Lys-rich leukemia (ELL) family protein, and an ELL-associated factor (EAF) protein (35). To circumvent this regulatory step, HIV-1 encodes a trans-activator protein (Tat) that binds to and recruits P-TEFb specifically to sites of nascent proviral transcription through recognition of a transactivation response (TAR) RNA stem loop produced at the immediate 3’ end of all viral RNA transcripts (36–38).

The distribution of and competition for P-TEFb binding among different complexes is an area of active investigation, with several strategies to enhance the biogenesis or availability of P-TEFb showing promise for HIV-1 latency reversal. For example, BET bromodomain inhibitors (such as JQ1) have been shown to be potent LRAs in *ex vivo* models by releasing P-TEFb from BRD4 (12). Likewise, the release of P-TEFb from the 7SK RNP complex has been shown to reactivate latent proviruses in *ex vivo* models (39). That being said, the release of P-TEFb from the 7SK RNP complex has been shown to directly correlate with increased BRD4 binding, suggesting that release from any one complex will not necessarily increase the amount of unbound P-TEFb or the amount recruited to specific sites of transcription (33). Furthermore, post-translational modification of P-TEFb (*i.e.*, through phosphorylation of CDK9 at Serine 175) has been shown to influence P-TEFb distribution in certain regulatory complexes, again highlighting the unique properties of release from each complex (40).

While the release of P-TEFb from the 7SK RNP complex and BET proteins such as BRD4 have been explored as strategies for HIV-1 latency reversal, the release of P-TEFb from SECs has not been explored. Previous work has demonstrated that HIV-1 Tat biochemically co-purifies with several SEC proteins (41, 42), though they seemingly have functionally redundant purposes in P-TEFb recruitment. In this study, we test the hypothesis that the SEC is not necessary for HIV-1 viral transcription and that the release of P-TEFb from cellular SEC complexes can serve as a novel strategy to reactivate latent HIV-1 proviruses.

## RESULTS

### The Super Elongation Complex is not required for HIV-1 replication in primary CD4+ T cells

While biochemical purification of HIV-1 Tat from cell lines has been shown to pull down other SEC members besides P-TEFb (including AFF1, AFF4, ELL2, ENL and AF9), the role of the SEC in HIV-1 replication in primary CD4+ T cells is unclear. To determine whether the SEC is required for HIV-1 replication in primary CD4+ T cells, we used multiplexed CRISPR-Cas9 ribonucleoproteins (RNPs) to knock-out expression of each SEC component in cells from multiple healthy donors (**Figure 1A**) (43–45). Each multiplexed reaction consisted of 4 or 5 independent guide RNA targeting the same gene (45). A non-targeting (NT) guide RNA was used as a negative control whereas a previously validated guide RNA targeting the HIV-1 co-receptor gene, *CXCR4*, was used as a positive control (43). Immunoblots of protein lysates collected 72 hours after CRISPR-Cas9 RNP electroporation demonstrated depletion of each target (**Figure 1B**). Visualization of AFF1 and AFF4 depletion required CCNT1 immunoprecipitation, likely due to their low steady-state levels in CD4+ T cells and low-affinity antibodies (**Figure 1C**). Notably, the depletion of some targets had secondary effects on other complex members. For example, knock-out of *CCNT1* decreased CDK9 steady state levels and knock-out of *AFF1*, *AFF4*, and *AF9* each decreased ENL steady-state levels. None of the knockouts were found to decrease cell viability significantly at this time point (**Supplemental Figure 1A-B**).

**Figure 1.**
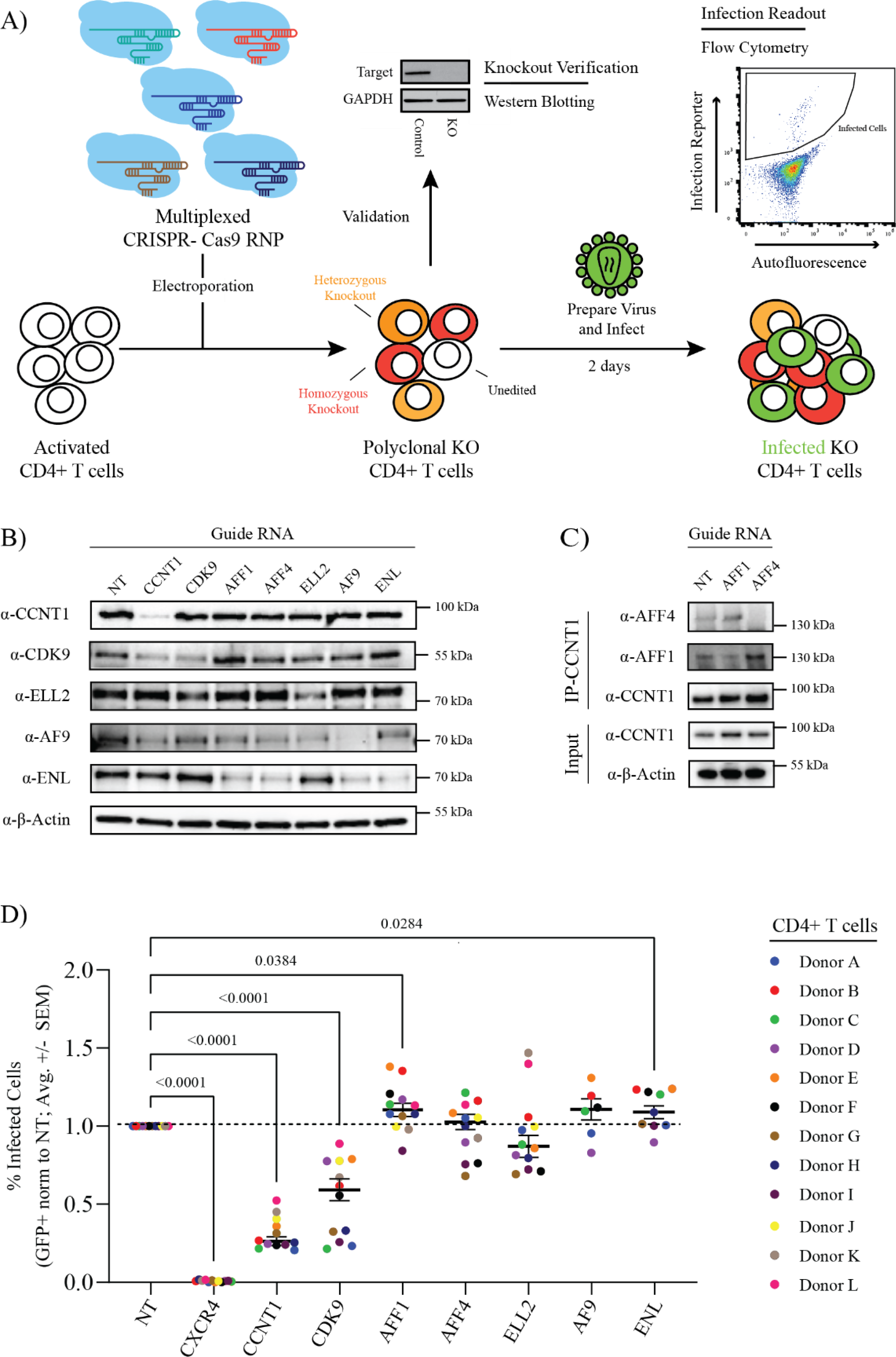
Knock-out of Super Elongation Complex members does not decrease HIV-1 infection in primary CD4+ T cells. **A)** Experimental schematic of multiplex CRISPR-Cas9 gene editing of primary CD4+ T cells. **B)** Immunoblots to assess CCNT1, CDK9, ELL2, AF9, and ENL knock-out efficiency in primary CD4+ T cells relative to a non-targeting (NT) control (1 representative donor). **C)** Immunoblots to assess AFF1 and AFF4 knock-out efficiency in primary CD4+ T cells relative to a NT control following immunoprecipitation of CCNT1 (1 representative donor). **D)** Percent infected (GFP+) primary CD4+ T cells (normalized to the donor-matched NT control) 48 hours after challenge with HIV-1 NL4.3 Nef-IRES-GFP. Each dot represents the average of technical triplicates; the black line represents the mean of means ± standard error of means. n = 12 donors for NT, CXCR4, CCNT1, CDK9, AFF1, AFF4, and ELL2; n=6 donors for AF9; and n=9 donors for ENL. Statistics were calculated by 2-way ANOVA with Dunnet’s Multiple Comparison Test; significant p-values (p < 0.05) are shown.

To determine the impact of SEC component knock-out on HIV-1 replication, we challenged each cell population with replication-competent HIV-1 NL4.3 containing an IRES-driven GFP reporter inserted after Nef (HIV-1 NL4.3 Nef:IRES:GFP) in technical triplicate. The percentage of infected cells was quantified at two-days post-challenge by flow cytometry (**Supplemental Figure 1C**) and normalized to the NT control (**Figure 1D**). Knock-out of *CXCR4* ablated CXCR4-tropic NL4.3 strain infection as expected. Knock-out of each P-TEFb component (*CCNT1*, *CDK9*) significantly decreased infection. However, knock-out of the other SEC components either did not alter HIV-1 infection (*AFF4*, *ELL2*, *AF9*) or increase infection (*AFF1*, *ENL*) significantly. These data suggest that, while P-TEFb is required, the rest of the SEC is dispensable for HIV-1 replication in primary CD4+ T cells.

### The SEC inhibitor KL-2 enhances HIV-1 infection in primary CD4+ T cells

Seeing that knock-out of several SEC components either had no effect or significantly increased infection of primary CD4+ T cells, we next wanted to determine the impact of chemical perturbation of this complex using the previously described SEC inhibitor, KL-2 (32, 46). KL-2 inhibits SEC function by disrupting the interaction between CCNT1 and AFF1/AFF4 without altering overall protein levels of P-TEFb (**Figure 2A**) (32). Given that the SEC is dispensable for HIV-1 replication, we hypothesized that KL-2 treatment may increase the bioavailability of P-TEFb, and increase infection. Activated CD4+ T cells from three independent donors were treated over a range of KL-2 concentrations for 24 hours and then challenged with HIV-1 NL4.3 Nef:IRES:GFP for 48 hours in technical triplicate. Higher concentrations of KL-2 dramatically decreased cell viability, with 3.125 μM being the highest tolerated dose with minimal toxicity (**Figure 2B**). This dose was sufficient to inhibit the CCNT1:AFF4 interaction in primary CD4+ T cells within 24 hours as determined by CCNT1 immunoprecipitation (**Figure 2C**). Compared to DMSO treated control cells, KL-2 treatment resulted in a dose-dependent increase in infection with significant increases observed at 1.56 μM and 3.125 μM (**Figure 2D**).

**Figure 2.**
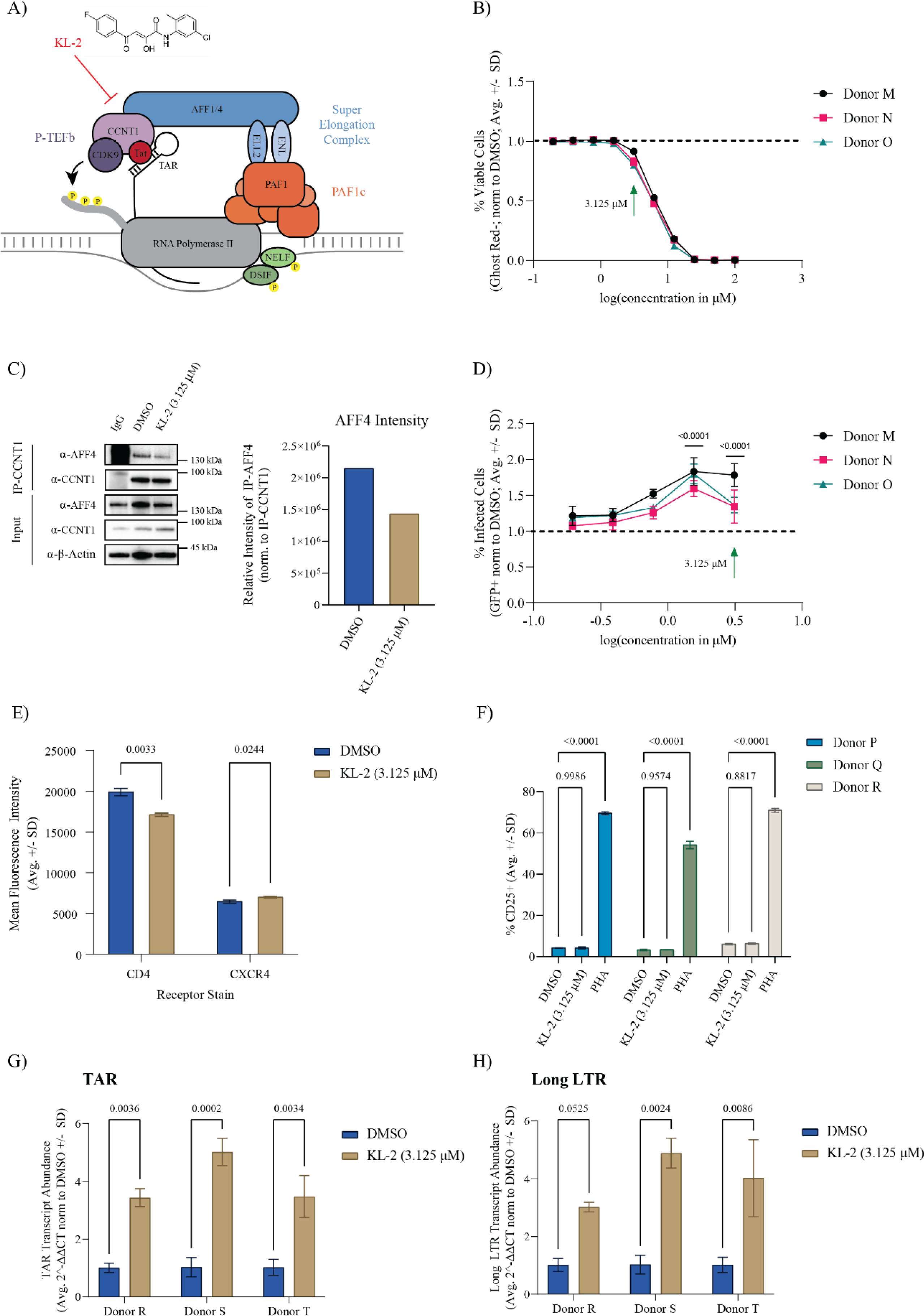
Super Elongation Complex disruptor KL-2 increases HIV-1 infection in primary CD4+ T cells. **A)** Model of Tat-mediated recruitment of P-TEFb to paused RNA Pol II at sites of nascent HIV-1 proviral transcription. **B)** Percent viable primary human CD4+ T cells (normalized to the donor-matched DMSO control) after 72 hours of treatment with increasing concentrations of KL-2 as measured by amine dye staining and flow cytometry. Data represent the average ± SD of technical triplicates; the green arrow indicates the concentration chosen for downstream experiments in primary CD4+ T cells. **C)** Immunoblots for AFF4 and CCNT1 following CCNT1 immunoprecipitation from primary CD4+ T cells treated for 24-hours with either DMSO or 3.125 μM KL-2 (1 representative donor). Quantification of immunoprecipitated AFF4 levels normalized to immunoprecipitated CCNT1 is shown on the right. **D)** Percent infected (GFP+) primary CD4+ T cells (normalized to the donor-matched DMSO control) 48 hours after challenge with HIV-1 NL4.3 Nef-IRES-GFP in the presence of increasing concentrations of KL-2 (24 hours pre-treatment before challenge). Data represent the average ± SD of technical triplicates (n = 3 donors); statistics were calculated relative to the DMSO control by two-way ANOVA and Sidak’s Multiple Comparison test with significant p-values (p < 0.05) shown. **E)** Percent of activated (CD25+) primary CD4+ T cells following treatment with DMSO, 3.125 μM KL-2, or 1μg/mL PHA for 48 hours as measured by immunostaining and flow cytometry. Data represent the average ± SD of technical triplicates (n = 3 donors); statistics were calculated by two-way ANOVA and Sidak’s Multiple Comparison test with p-values shown. **F)** Mean fluorescence intensity (MFI) of CD4 and CXCR4 on primary CD4+ T cells following treatment with DMSO or 3.125 μM KL-2 for 48 hours as measured by immunostaining and flow cytometry. Data represent the average ± SD of technical triplicates (n = 1 donor); statistics were calculated by Student’s t-test with p-values shown. **G)** Relative transcript levels of HIV-1 TAR and **H)** long LTR to human b-Actin in activated primary CD4+ T cells after 48-hours of challenge with HIV-1 in the presence or absence of a 24-hour pretreatment with KL-2. Data represent the mean of means of 3 biological replicates in technical duplicate ± standard error; statistics were calculated by two-way ANOVA with Sidak’s Multiple Comparison test.

Given that receptor and co-receptor expression can alter HIV-1 susceptibility, we next tested the impact of KL-2 treatment on CD4 and CXCR4 cell surface expression. Activated, primary CD4+ T cells from one representative donor were treated with 3.125 μM KL-2 or DMSO for 48 hours prior to immunostaining and flow cytometry (**Supplemental Figure 2A-B**). KL-2 treatment resulted in a slight, but significant increase in CXCR4 expression (mean fluorescence intensity) and a slight, but significant decrease in CD4 expression (**Figure 2E**). To determine if KL-2 could alternately impact CD4+ T cell activation, we treated unstimulated CD4+ T cells from three independent donors with DMSO, 3.125 μM KL-2, or the T cell mitogen Phytohemagglutinin (PHA) for 48 hours. Unlike PHA, which resulted in robust activation, KL-2 treatment did not impact T cell activation as measured by CD25 cell surface staining (**Figure 2F**). These results suggest that increased infection in the presence of KL-2 is not likely driven by changes in receptor expression.

To assess whether this increase in infection was due to enhanced viral transcription, activated CD4+ T cells from three healthy donors were treated with 3.125 μM KL-2 or DMSO for 24 hours and then challenged with HIV-1 NL4.3 Nef:IRES:GFP in technical triplicate. After 48 hours, RNA from the infected cultures was isolated and the expression of viral transcripts was measured using quantitative reverse transcription PCR (qRT-PCR). Quantification of TAR and long LTR transcripts were used to measure viral transcriptional initiation and elongation, respectively, relative to the human housekeeping gene, *β-Actin*. We found that KL-2 treatment significantly increased the expression of both TAR and long LTR transcripts (**Figure 2G-H**). These data support the genetic data above that the interaction between P-TEFb and the larger SEC is not required for HIV-1 replication in primary CD4+ T cells and that SEC disruption can enhance viral transcription.

### KL-2 synergizes with other latency reversing agents to reactivate latent infection in J-Lat models

The release of P-TEFb from sequestration by BRD4 or the 7SK complex has proven effective in latency reactivation (39, 47). Given that the SEC is not required for HIV-1 replication, we hypothesized that releasing P-TEFb from cellular SECs may also reactivate latent proviruses. To test this hypothesis, we treated J-Lat 5a8 cells (a Jurkat subclone harboring a silent, integrated non-replicative full-length provirus and a GFP reporter gene (48)) over a range of KL-2 concentrations for 48 hours. As in primary CD4+ T cells, higher concentrations of KL-2 decreased J-Lat 5A8 cell viability (**Figure 3A**). However, treatment with KL-2 alone did not significantly increase proviral reactivation (GFP+ cells as measured by flow cytometry) until 6.25 μM, at which dose the cells are only about 50% viable (**Figure 3A-B**). This dose was sufficient to inhibit the CCNT1:AFF4 interaction in J-Lat 5A8 cells within 24 hours, as determined by CCNT1 immunoprecipitation (**Figure 3C**).

**Figure 3.**
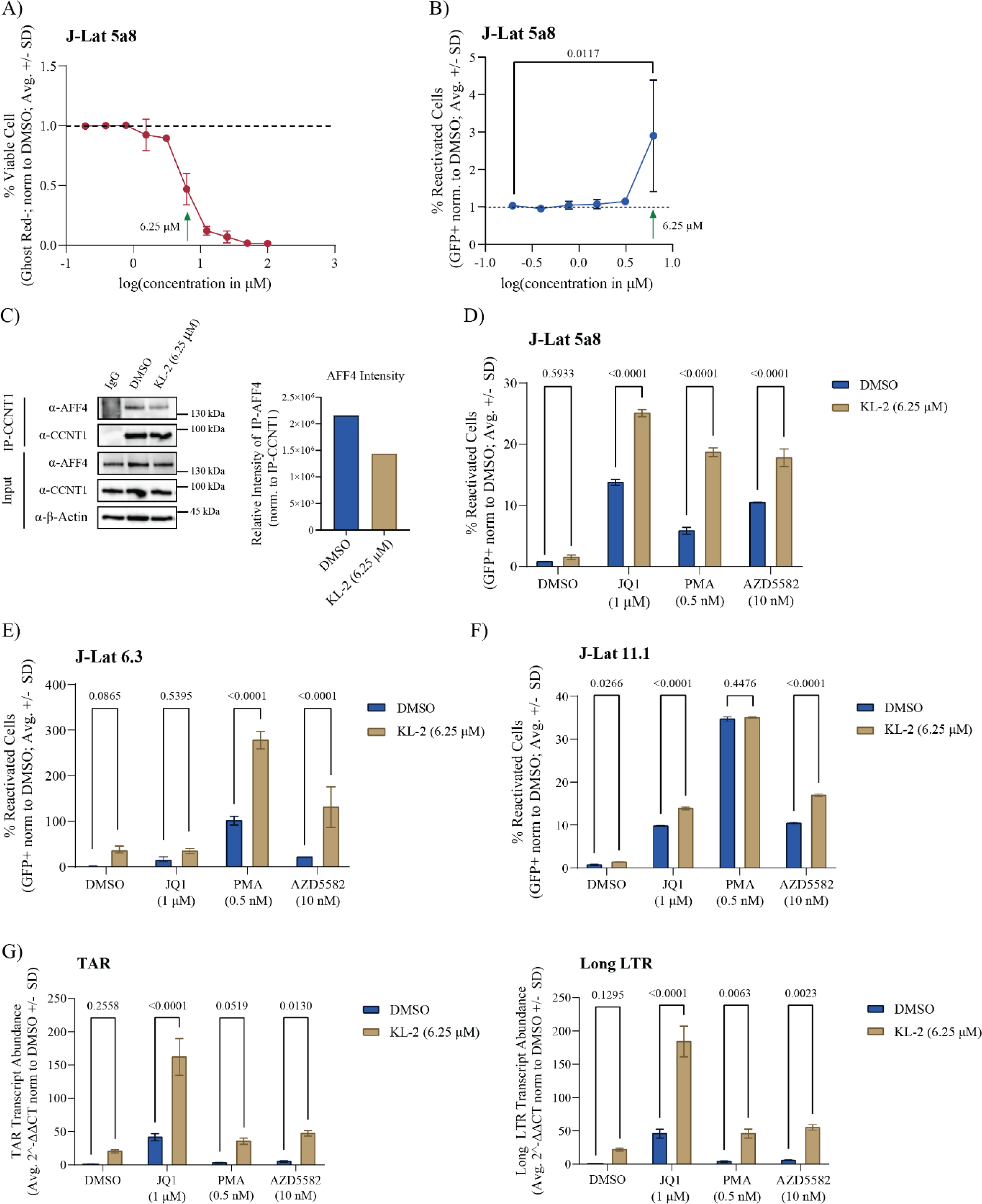
KL-2 enhances latency reversing agent activity in J-Lat cell lines. **A)** Percent viable J-Lat 5A8 cells (normalized to the DMSO control) after 48 hours of treatment with increasing concentrations of KL-2 as measured by amine dye staining and flow cytometry. Data represent the average ± SD of technical triplicates; the green arrow indicates the concentration chosen for downstream experiments in cell lines. **B)** Percent reactivated (GFP+) J-Lat 5A8 cells (normalized to the DMSO control) after 48 hours of treatment with increasing concentrations of KL-2. Data represent the average ± SD of technical triplicates; statistics were calculated relative to the DMSO control by two-way ANOVA and Sidak’s Multiple Comparison test. **C)** Immunoblots for AFF4 and CCNT1 following CCNT1 immunoprecipitation from J-Lat 5A8 cells treated for 24 hours with either DMSO or 6.25 μM KL-2. Quantification of immunoprecipitated AFF4 levels normalized to immunoprecipitated CCNT1 is shown on the right. **D)** Percent reactivated (GFP+) J-Lat 5A8 cells, **E)** J-Lat 6.3 cells, or **F)** J-Lat 11.1 cells (normalized to the DMSO control) after 48 hours of treatment with the indicated compounds in the presence or absence of KL-2. Data represent the average ± SD of technical triplicates; statistics were calculated by two-way ANOVA with Sidak’s Multiple Comparison test. **G)** Relative transcript levels of HIV-1 TAR (left) and long LTR (right) to human b-Actin in J-Lat 5A8 cells after 48 hours of treatment with the indicated compounds in the presence or absence of KL-2 (normalized to the DMSO control). Data represent the mean of means of 3 biological replicates in technical duplicate ± standard error; statistics were calculated by two-way ANOVA with Sidak’s Multiple Comparison test.

While KL-2 was not sufficient to reactivate latent proviruses in J-Lat 5A8 cells, we hypothesized that it could enhance the activity of other latency reversing agents (LRAs) that act through different mechanisms, similar to the PAF1 complex inhibitors we reported previously (16). To test this hypothesis, we treated J-Lat 5a8 cells with several well-characterized LRAs in the presence or absence of KL-2, including JQ1 (a BRD4 inhibitor that enhances P-TEFb availability and relieves chromosomal repression), Phorbol 12-myristate 13-acetate (PMA, a protein kinase C activator that activates canonical NFkB transcription), and AZD5582 (an IAP antagonist that activates non-canonical NFkB transcription). While KL-2 alone did not significantly increase reactivation compared to the DMSO control, it significantly enhanced reactivation in the presence of the other three LRAs (**Figure 3D**). This same result was observed in two other J-Lat clones with different proviral integration sites: J-Lat 6.3 cells (**Figure 3E**) and J-Lat 11.1 cells (**Figure 3F**). Paired cell viability data is provided in **Supplemental Figure 3A-C.**

To assess whether the effect of KL-2 in combination with other LRAs was synergistic, we calculated excess over Bliss scores (49) for KL-2 in combination with JQ1 and AZD5582. Excess over Bliss calculations measure whether the observed combinatorial effects at given concentrations are above or below predicted additivity, where a score of 0 is additive, scores above 0 are considered synergistic, and scores less than 0 are considered antagonistic. At the fixed 6.25 μM concentration of KL-2 used in the previous assays in combination with ascending doses of AZD5582, we observed positive excess over Bliss scores ranging from 0.46 to 0.94, suggestive of synergistic effects (**Supplemental Figure 3D**; top). Notably, at a fixed concentration of AZD5582 (10 nM), we observe strictly additive effects of KL-2 until the effective concentration of 6.25 μM is reached. Similar trends are observed with JQ1. At the fixed 6.25 μM concentration of KL-2 with ascending doses of JQ1, we again saw positive excess over Bliss scores ranging from 0.54 to 0.94 (**Supplemental Figure 3D**; bottom). Likewise, at a fixed concentration of JQ1 (1 μM), we observe strictly additive effects of KL-2 until the effective concentration of 6.25 μM is reached. This data would indicate that at concentrations of KL-2 lower than 6.25 μM we are likely not seeing optimal disruption of the SEC, but upon disruption at the effective dose of KL-2, the release of P-TEFb acts synergistically with the LRAs tested.

Given the role of KL-2 in P-TEFb release from the SEC, we hypothesized that treatment with KL-2 would enhance transcriptional elongation. To test this, we repeated our combinatorial LRA treatments with and without KL-2 in J-Lat 5A8 cells and extracted RNA at 48 hours post-treatment to quantify viral transcripts using qRT-PCR. HIV-1 long LTR transcripts, made after the start of transcriptional elongation, were not statistically increased upon KL-2 treatment alone, but were enhanced by KL-2 addition to every other tested LRA (**Figure 3G**, right). HIV-1 TAR transcripts, made immediately after transcriptional initiation, were not statistically increased upon KL-2 addition either alone or in the presence of PMA, however, KL-2 treatment did significantly increase transcriptional initiation in the presence of JQ1 and AZD5582 (**Figure 3G**, left). Taken together, these results suggest that KL-2 can enhance the latency reactivation activity of other LRAs in J-Lat models by increasing the availability of P-TEFb and promoting transcriptional elongation. However, additional impacts on transcriptional initiation cannot be ruled out as also observed in our primary CD4+ T cell data (**Figure 2G**).

### Assessing the Tat dependency of KL-2 latency reversal

The SEC is dispensable for HIV-1 replication in primary CD4+ T cells, and release of P-TEFb from the SEC promotes latency reversal in J-Lat cells, suggesting that P-TEFb is recruited to paused RNA Pol II at proviral integration sites in an SEC-independent manner in these models. This is most likely through the viral accessory protein Tat, which can recruit P-TEFb directly to sites of nascent viral transcription through an interaction with the TAR stem-loop on the 5’ end of viral transcripts (though alternate mechanisms for P-TEFb recruitment have been described). To assess the Tat-dependency of latency reactivation in the J-Lat 5A8 cell line model, we performed combinatorial treatments with a panel of LRAs in the presence and absence of KL-2 and in the presence of two different Tat inhibitors: Spironolactone and Triptolide (**Figure 4A**; viability data in **Supplemental Figure 4A**). Spironolactone induces the degradation of the XBP helicase, a component of the TFIIH initiation complex, which inhibits Tat-dependent transcription (50), while Triptolide promotes proteasomal degradation of the Tat protein itself (51). In the presence of the DMSO control, KL-2 enhanced the latency reactivation activity of JQ1, PMA, and AZD5582 as observed previously (**Figure 3D**). Both Tat inhibitors dramatically reduced the reactivation potency of each LRA, in some cases to near baseline levels. Treatment with KL-2, however, still enhanced reactivation in combination with JQ1 and PMA, even though KL-2 treatment alone had no effect. These data suggest that either there is a residual amount of Tat or an SEC/Tat-independent route for P-TEFb recruitment to sites of proviral transcription when P-TEFb is released from the SEC. Notably, KL-2 did not enhance the reactivation potential of AZD5582 upon Tat inhibition, suggesting that this LRA may be uniquely dependent on Tat and/or the SEC for P-TEFb recruitment (**Figure 4A**).

**Figure 4.**
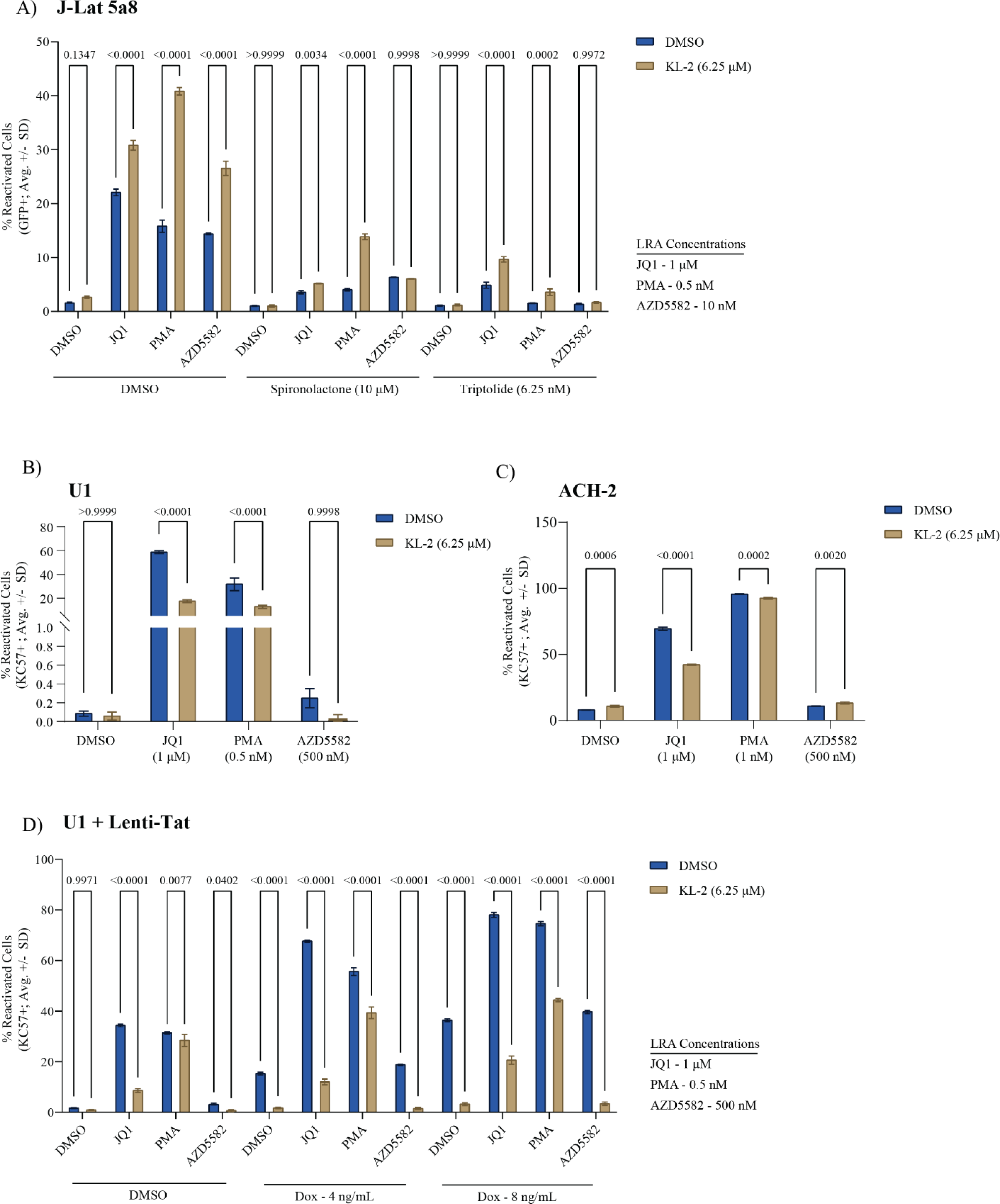
KL-2 latency reversing activity is Tat- and cell line-dependent. **A)** Percent reactivated (GFP+) J-Lat 5a8 cells after 48 hours of treatment with indicated compounds in the presence or absence of KL-2 and the Tat inhibitors Spironolactone and Triptolide. **B)** Percent reactivated (KC57-FITC+) U1 cells after 48 hours of treatment with indicated compounds in the presence or absence of KL-2. **C)** Percent reactivated (KC57-FITC+) ACH-2 cells after 48 hours of treatment with indicated compounds in the presence or absence of KL-2. **D)** Percent reactivated (KC57-FITC+) Lenti-Tat U1 cells after 48 hours of treatment with indicated compounds in the presence or absence of KL-2 and with Tat induction at three different concentrations of doxycycline (0, 4, and 8 ng/mL). For all panels, the data represent the average ± SD of technical triplicates; statistics were calculated by two-way ANOVA with Sidak’s Multiple Comparison test.

To exclude the potential effects of residual amounts of Tat, we next tested the impact of SEC disruption in two cell line models of latency that lack a functional Tat-TAR axis: U1 cells and ACH-2 cells. The U1 cell line is a U937-based monocytic cell line that harbors two copies of the HIV-1 provirus, one of which expresses no Tat due to the lack of a start codon and the second of which encodes a Tat mutant with suboptimal P-TEFb affinity (52). The ACH-2 cell line is a T cell line with one integrated provirus that encodes a fully functional Tat, but that lacks a functional TAR stem loop (53). Both cell lines were treated with the same LRA panels as above in the presence and absence of KL-2 for 48 hours. These cell lines lack a fluorescent reporter; therefore, reactivation was monitored by intracellular p24 immunostaining and flow cytometry (viability data in **Supplemental Figure 4B-C)**. Treatment with KL-2 alone caused negligible reactivation in either cell type, like the J-Lat models (**Figure 4B-C**). While JQ1 and PMA treatment resulted in strong reactivation, AZD5582 had a minimal effect, consistent with the aforementioned dependency on Tat. In contrast with the J-Lat models, however, combinatorial treatment with KL-2 significantly decreased the reactivation potential of JQ1 and PMA in both cell lines, suggesting that reactivation by these LRAs is at least in part SEC-dependent (**Figure 4B-C**).

These data suggest that while the SEC may be dispensable for Tat-dependent transcription and reactivation, it may play an essential role in P-TEFb recruitment without functional Tat. If so, the delivery of exogenous Tat to U1 cells might be sufficient to circumvent the need for the SEC, reversing the KL-2 treatment phenotype. To test this hypothesis, we transduced U1 cells with a doxycycline (Dox)-inducible lentiviral construct encoding full-length HIV-1 NL4.3 Tat (referred to as Lenti-Tat), selecting for a pure polyclonal population of transduced cells in puromycin. The U1 + Lenti-Tat cells were treated with a range of Dox concentrations, resulting in a dose-dependent increase in proviral reactivation (**Supplemental Figure 4D**). U1 + Lenti-Tat cells were then treated with the same panel of LRAs in the presence or absence of KL-2 in the presence of either 0, 4, or 8 ng/mL Dox (**Figure 4D**; viability data in **Supplemental Figure 4E**). Without Tat induction, KL-2 suppressed JQ1- and PMA-mediated reactivation as before (**Figure 4B**). Tat induction by Dox was sufficient to induce a basal level of reactivation, which was almost entirely reversed by KL-2 treatment (**Figure 4D**). JQ1 and PMA induced reactivation above the Tat-induced baseline, but retained sensitivity to KL-2, while AZD5582 failed to reactivate above baseline even in the presence of Tat. These data suggest that the presence of Tat alone is not sufficient to dictate the route of P-TEFb recruitment to paused RNA Pol II, and that the SEC may play an essential role in P-TEFb recruitment in some cell types or at some integration sites.

### KL-2 increases HIV-1 viral transcripts in PBMCs from aviremic patients

Blocks to transcriptional elongation have been shown to contribute to HIV-1 latency maintenance in cells from virally suppressed people living with HIV (PLWH) (13, 14). The release of P-TEFb sequestration from BRD4 using bromodomain inhibitors (such as JQ1) has proven effective at reactivating viral gene expression in peripheral blood mononuclear cells (PBMCs) from these patients (54). We hypothesized that KL-2 would likewise reactivate viral gene expression in patient PBMCs. To test this, cryopreserved PBMCs were obtained from five PLWH enrolled in the Northwestern University Clinical Research Site for the MACS/WIHS Combined Cohort Study (MWCCS) (**Figure 5A**). The selected PLWH had been virally suppressed with ART for more than five years with undetectable HIV-1 plasma levels at the time of blood draw (<50 copies/ml) (**Figure 5B**). Cells from these patients were treated for 48 hours with DMSO, JQ1, or AZD5582 alone or in combination with KL-2. RNA was isolated from the treated cells and reactivation of viral gene expression was measured by qRT-PCR for HIV-1 *gag* (relative to the human housekeeping gene *LDHA*, **Figure 5C**).

**Figure 5.**
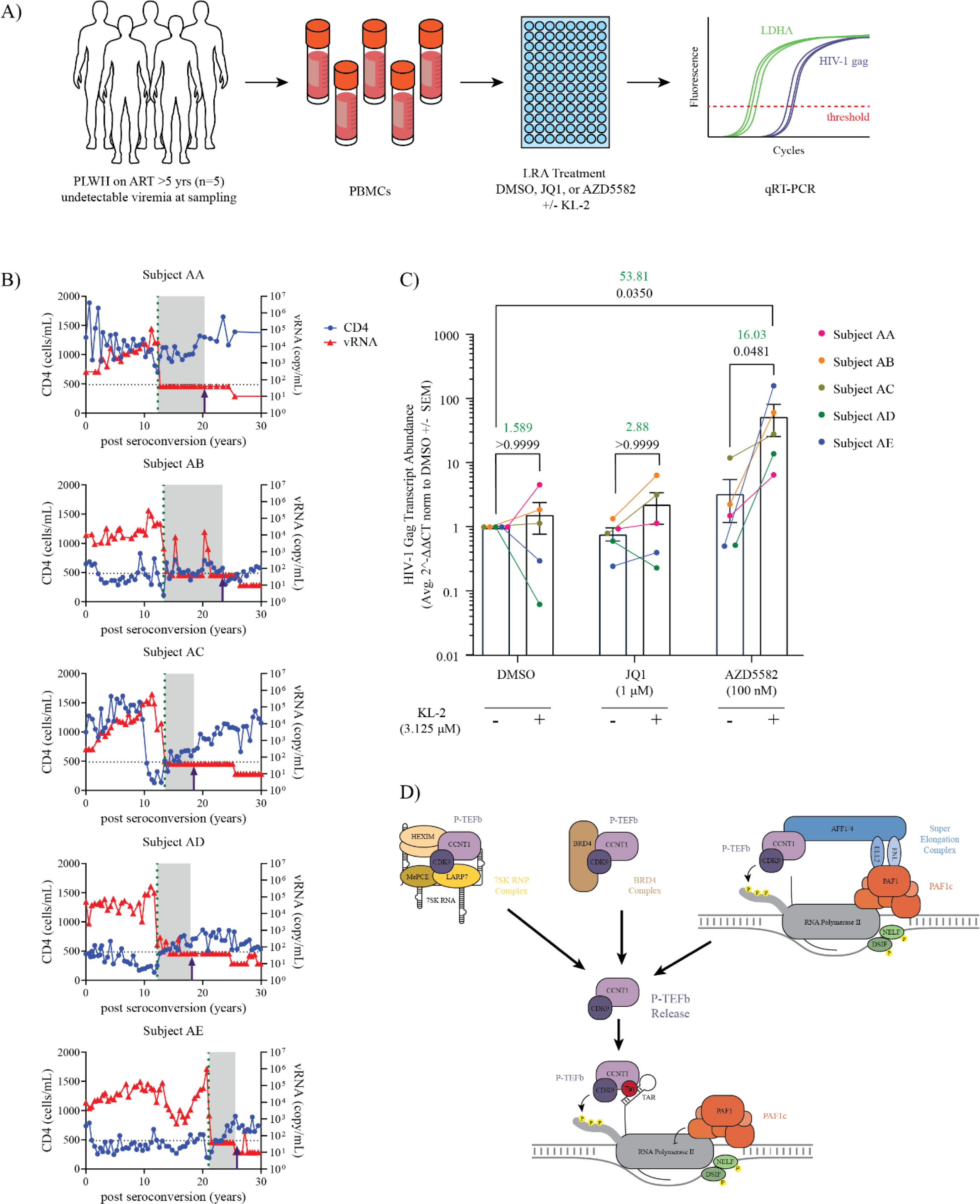
KL-2 enhances latency reversing agent activity in PBMCs from virally suppressed people living with HIV. **A)** Experimental schematic of the latency reversal assay using PBMCs from five HIV-1 patients with undetectable viral loads. Intracellular *gag* transcript levels were measured by qRT-PCR after treatment with JQ1, PMA, or AZD5582 in the presence or absence of KL-2. **B)** Overlaid line graphs depicting the CD4 count (cells/mL, blue) and levels of viral RNA (copies/mL, red) in each subject over time. The dotted line shows the time of antiretroviral therapy initiation, and the time of the blood draw used for this assay is depicted by an arrow. **C)** Relative transcript levels of HIV-1 *gag* relative to human *LDHA* in patient PBMCs (n = 5 donors) after 48 hours of treatment with the indicated compounds in the presence or absence of KL-2. Data represent the average 2-ΔΔCt [(Ct_gag_-Ct_LDHA_)_LRAs_ - ( Ct_gag_-Ct_LDHA_)_DMSO_] ± SEM of technical triplicates. Statistics were calculated using a two-way ANOVA with Sidak’s Multiple Comparison Test. The fold change in mean 2-ΔΔCt is shown above for relevant comparisons in green text with corresponding p-values shown below in black text. **D)** Model of cellular P-TEFb reservoirs whose disruption have demonstrated activity in HIV-1 latency reversal.

KL-2 alone resulted in a slight increase in *gag* expression in three out of the five donors with the mean transcript levels increasing 1.5-fold, although this increase was not statistically significant (**Figure 5C**). Surprisingly, treatment with JQ1 alone was not sufficient to induce *gag* expression in these donors, though the addition of JQ1 and KL-2 resulted in a 2.88-fold increase over JQ1 treatment alone (not significant, **Figure 5C**). Likewise, AZD5582 treatment had minimal impact on *gag* expression alone. However, consistent with our results in cell line models, we observed that KL-2 significantly enhanced the ability of AZD5582 to reactivate HIV-1 transcription across all donors with a marked 53-fold increase over the DMSO only control and a 16-fold increase compared to AZD5582 alone (**Figure 5C**). Taken together, these data demonstrate that the release of P-TEFb from cellular SECs using the small molecule KL-2 can enhance the effects of a range of LRAs in cell line models of HIV-1 latency as well as improve transcriptional reactivation by AZD5582 in primary PBMCs from HIV-1 patients.

## DISCUSSION

In this study, we demonstrate a potential new strategy for enhancing HIV-1 latency reversal through the release of P-TEFb from the cellular pool of SECs. We show that KL-2, a small molecule inhibitor of the interaction between the SEC and P-TEFb, is sufficient to enhance viral transcription in primary CD4+ T cells and can synergistically enhance the activity of other LRAs in cell line models of latency. Finally, we demonstrate that KL-2 can increase HIV-1 *gag* expression in PBMCs from PLWH on suppressive ART, most notably in combination with the non-canonical NFkB agonist, AZD5582. We propose a model in which KL-2 release of P-TEFb from the cellular pool of SECs enhances Tat-dependent transcriptional elongation of integrated proviruses, akin to BET bromodomain inhibitors and 7SK RNP inhibitors (**Figure 5D**). These results have several implications for our understanding of viral transcription and future directions. Latency reversal through the release of P-TEFb from cellular SECs has yet to be previously explored likely due to the perceived dependency of viral transcription on the SEC. This effect has been supported by biochemical purifications of HIV-1 Tat from human cell lines that revealed interactions with a larger SEC (41, 42), as well as by genetic knock-down experiments in cell lines showing a decrease in Tat-dependent transcription upon SEC component depletion (55, 56). However, given that both the SEC and Tat recruit P-TEFb to sites of nascent transcription, they share some functional redundancy. We found that knock-out of SEC components from activated, primary CD4+ T cells from 12 independent donors did not inhibit viral replication, suggesting that the SEC is not required in this cellular context. This result independently verified using KL-2, which inhibits the interaction between CCNT1 (P-TEFb) and AFF1/4 (of the larger SEC).

While we expected KL-2 to enhance the transcriptional elongation of integrated proviruses due to the release of P-TEFb from cellular SECs, we also saw increases in transcriptional initiation as measured by qRT-PCR for TAR transcript levels. Likewise, in J-Lat 5A8 cells, we saw increases in transcriptional elongation and initiation when KL-2 was added with other LRAs, even though KL-2 addition alone was not sufficient for reactivation as measured by GFP positivity or by viral transcript. This finding suggests that either KL-2 has secondary effects not mediated by P-TEFb or that the redistribution of P-TEFb away from SECs has secondary effects that could impact transcriptional initiation at proviral integration sites. BET bromodomain inhibitors have also been reported to increase HIV-1 transcriptional initiation (12, 13), though these effects have been suggested to occur through modulation of the epigenetic regulatory functions of these proteins (57). Other reports have indicated that P-TEFb mediated release of paused RNA Pol II can result in enhanced transcriptional initiation simply by increasing the number of transcribing polymerases (58). Still, this connection between transcriptional elongation and initiation has yet to be fully understood in the context of HIV-1 transcription.

While KL-2 was sufficient to boost viral replication in activated, primary CD4+ T cells, in and of itself it displayed minimal reactivation potential in both cell line models of latency and in PBMCs from PLWH on suppressive ART. This finding is similar to our recent report of a novel inhibitor of the PAF1 complex (iPAF1C) that had minimal activity on its own, but greatly enhanced the reactivation potential of other LRAs (16). Both cases highlight the multifaceted nature of the blocks to viral gene expression that underlie the latent state as well as the limitations to single agent drug screening to identify promising, next-generation LRAs. Combinatorial approaches to dissect the genetic underpinnings of HIV-1 latency and discover new, synergistic drug interactions should be prioritized.

While KL-2 alone failed to significantly increase HIV-1 *gag* transcript levels in patient PBMCs, in combination with AZD5582 it resulted in a 16-fold increase over AZD5582 treatment alone and a 53-fold increase over the DMSO control. Crosswise dose titrations in J-Lat 5A8 cells showed a strong synergistic potential between AZD5582 and KL-2. This is consistent with reports of robust synergy between AZD5582 and P-TEFb release through BET bromodomain inhibition (12). Notably, the BET bromodomain inhibitor JQ1 showed minimal reactivation activity in our patient PBMCs, even in the presence of KL-2, in contrast to our cell line data. This finding could reflect stochastic differences driven by variations in patient characteristics, integration site, chromatin state, transcription factor availability, etc (13). Future work will compare P-TEFb release from SECs to release from other cellular reservoirs, such as BRD4 or the 7SK RNP. Recent studies have shown that post-translational modifications of P-TEFb, most notably phosphorylation of CDK9 Serine 175 and Threonine 186, can drive inclusion into different complexes and may strongly influence bioavailability and activity (59–62). Therefore, it is possible that disruption of complexes housing ‘active’ P-TEFb is a more direct route to redirecting P-TEFb activity.

This is not to say that the SEC is never required for viral transcription. In latency model cell lines that lacked a functional Tat (U1 cells) or TAR stem loop (ACH-2 cells), KL-2 inhibited the reactivation potential of several LRAs, suggesting that viral transcription may be more dependent on the SEC when Tat is either defective or not expressed. We first attempted to test this by inhibiting Tat in the J-Lat 5A8 model cell line using two previously described Tat inhibitors, Triptolide and Spironolactone. While both compounds reduced LRA efficacy, KL-2 still boosted the activity of JQ1 and PMA, but not AZD5582. This suggests that P-TEFb can be recruited to proviral integration sites in a Tat and SEC-independent manner upon PMA or JQ1 treatment, potentially through a transcription factor such as NFkB. The inability of AZD5582 to do so suggests that non-canonical NF-kB activation does not recruit the same milieu of transcription factors, making it uniquely Tat or SEC dependent. This is consistent with the complete lack of activity of AZD5582 in cell line models lacking functional Tat/TAR activity and may underlie the remarkable synergy between non-canonical NFkB agonists and agents that release P-TEFb (12). Additionally, triptolide has been characterized outside of HIV-1 transcription in its ability to prevent RNA Pol II reinitiation following pausing through inhibition of xeoderma pigmentosum group B-complementing protein (XPB) and is often used as a tool compound for measuring the fate of paused RNA Pol II at different time points (63, 64). With this in mind, it is possible that compounds JQ1 and PMA result in *de novo* recruitment of RNA Pol II thereby increasing transcriptional initiation whereas AZD5582 may be more reliant on RNA Pol II pause-release.

To further explore the Tat dependency of KL-2, we tried to rescue Tat function in the U1 cell line using a Dox-inducible system. We hypothesized that by providing Tat, the SEC would no longer be required for viral gene expression such that SEC disruption by KL-2 would enhance reactivation as seen in the J-Lat models. While Tat induction itself was sufficient for reactivation, this reactivation was completely abolished by the addition of KL-2. Even when Tat was minimally induced and other LRAs were added, KL-2 still inhibited reactivation, suggesting that additional factors - such as steady-state levels of P-TEFb, integration site, epigenetic factors, or transcription factor availability - may drive SEC dependency besides just the presence or absence of Tat. Indeed, SEC disruption by KL-2 in the original report of the inhibitor demonstrated an outsized impact on Myc-dependent transcription (32), suggesting that additional factors driving the SEC dependency of proviral transcription have yet to be described. Regardless, the dual-acting nature of KL-2 in enhancing latency reactivation in some circumstances (*i.e.*, when Tat is present) and inhibiting latency reactivation in others (*i.e.*, when the cell state dictates SEC dependency) presents a unique opportunity to leverage the heterogenous nature of the latent reservoir to both reverse and promote latency.

Taken together, our results indicate that release of P-TEFb from cellular SECs is a novel mechanism for promoting HIV-1 viral transcription during both active and latent infection. We demonstrated the enhancement of latency reversal in multiple latent cell line models and in primary PBMCs from PLWH on suppressive ART. This work demonstrates the importance of increasing the production or availability of free P-TEFb for recruitment to viral loci as a powerful strategy for bolstering current LRAs, most notably non-canonical NF-kB agonists. Due to the heterogeneity of blocks to viral replication in the latent reservoir, it is likely that combinatorial LRA treatments will be the best strategy for potent latency reversal moving forward. Further efforts are needed to understand the intracellular distribution of active P-TEFb to characterize the most critical reservoir to target to enhance transcription. Additionally, our work demonstrates that disruption of SECs could enhance latency reversal or promote the maintenance of latency depending on the cellular context. Understanding the mechanism that controls this molecular switch would aid in understanding whether SEC disruptors could be a viable dual-acting molecule to aid in finding a functional cure for HIV-1 infection.

## METHODS

### CD4+ T cell isolation

Primary human CD4+ T cells from healthy donors were isolated from leukoreduction chambers after Trima apheresis (STEMCELL Technologies). PBMCs were isolated by Ficoll centrifugation. Bulk CD4+ T cells were subsequently isolated from PBMCs by magnetic negative selection using an EasySep Human CD4+ T cell isolation kit (STEMCELL Technologies; per the manufacturer’s instructions). Isolated CD4+ T cells were suspended in RPMI 1640 (Sigma-Aldrich) supplemented with 5 mM HEPES (Corning), penicillin-streptomycin (50 mg/ml; Corning), 5 mM sodium pyruvate (Corning), and 10% HI FBS (Gibco). Media were supplemented with interleukin-2 (IL-2; 20 IU/ml; Miltenyi) immediately before use. For activation, bulk CD4+ T cells were immediately plated on anti-CD3–coated plates coated for 2 hours at 37°C with anti-CD3 (20 mg/ml) (UCHT1; Tonbo Biosciences)] in the presence of soluble anti-CD28 (5 mg/ml; CD28.2; Tonbo Biosciences). Cells were stimulated for 72 hours at 37°C and 5% CO_2_ before treatment with KL-2.

### crRNP production

Lyophilized crRNA and tracrRNA (Dharmacon) was resuspended at a concentration of 160 µM in 10 mM Tris-HCL (7.4 pH) with 150 mM KCl. Cas9 ribonucleoproteins (RNPs) were made by incubating 5 µL of 160 µM crRNA (Horizon) with 5 µL of 160 µM tracrRNA for 30 minutes at 37 °C, followed by incubation of the gRNA:tracrRNA complex product with 10 µL of 40 µM Cas9 (UC Berkeley Macrolab) to form RNPs. Five 3.5 µL aliquots were frozen in Lo-Bind 96-well V-bottom plates (E&K Scientific) at −80 °C until use. For synthesis of multiplexed RNPs, four independent crRNA targeting the same gene were mixed at a 1:1:1:1 ratio prior to addition of the tracrRNA as above.

### CRISPR RNA Sequences

crRNA was ordered from Horizon Discovery using either the catalog numbers for predesigned guide sequences or custom guide sequences in the table below.

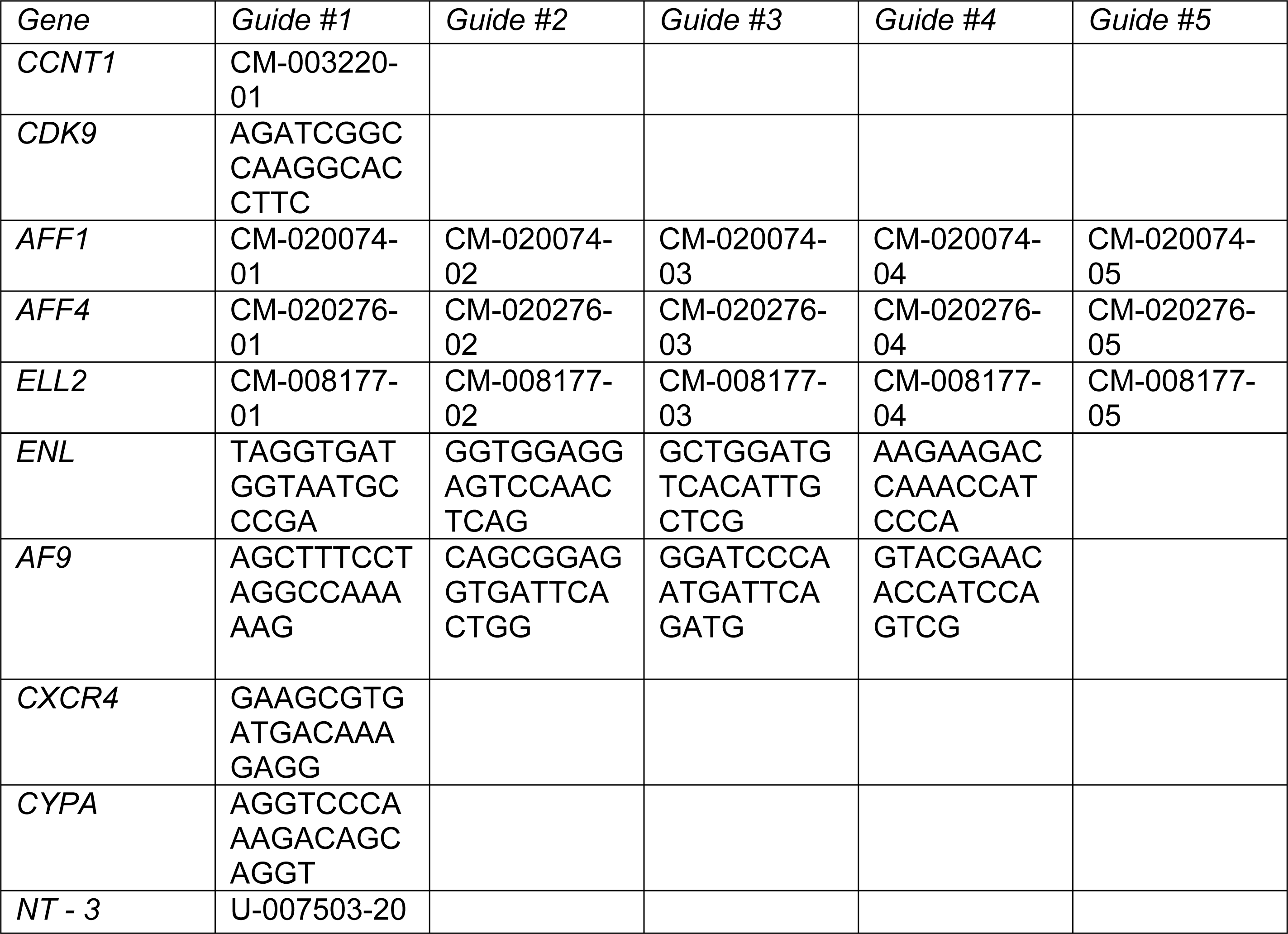

### CD4+ T cell CRISPR Knockouts

Following CD4+ T cell isolation and stimulation, as above, cells were counted, centrifuged at 400 × g for 5 minutes, the supernatant was removed by aspiration, and the pellet was resuspended in 20 µL of supplemented room-temperature P3 electroporation buffer (Lonza) per reaction. Each reaction consisted of 1 × 10^6^ cells, 3.5 µL of RNP, and 20 µL of electroporation buffer. The cell suspension was then gently mixed with thawed RNP and aliquoted into a 96-well electroporation cuvette for electroporation with the 4D 96-well shuttle unit (Lonza) using pulse code EH-115. Immediately after electroporation, 80 µL of prewarmed media without IL-2 was added to each well and cells were allowed to rest for at least one hour in a 37 °C cell culture incubator. Subsequently, cells were moved to 96-well flat-bottom culture plates prefilled with 100 µL of warm complete media with IL-2 at 40 IU/mL (for a final concentration of 20 IU/mL) and anti-CD3/anti-CD2/anti-CD28 beads (T cell Activation and Stimulation Kit, Miltenyi) at a 1:1 bead:cell ratio.

### Whole Cell Protein Lysate Preparation

Whole cell lysates were prepared by suspension of cell pellets (Typically ∼150,000 cells) directly in 2.5x Laemmli Sample Buffer followed by denaturization at 98°C for 30 minutes.

### Affinity Purification of endogenous CCNT1

10 million cells were pelleted, washed with PBS, and subsequently resuspended in 1 mL of lysis buffer (0.5% NP40, 50 mM Tris–HCl pH 7.4; 150 mM NaCl, 1 mM EDTA, cOmplete protease (Roche) and PhosSTOP phosphatase (Roche) inhibitors). Samples were then rotated at 4C for 30 minutes and transferred to -80°C for at least 30 minutes to complete cell lysis. Following lysis, samples were thawed on ice and centrifuged in a prechilled microcentrifuge at 3500 x g for 20 minutes to remove cellular debris. Lysates were then precleared by incubation for 1 hour at 4°C while rotating with 50 μL protein A agarose beads (Cell Signaling Technologies, Cat #9863) CCNT1 antibody (Cell Signaling Technologies, Cat#81464) was then added to precleared lysates at 1:100 and incubated with rotation overnight at 4°C. After incubation, 50 μL of fresh Protein A agarose beads were added to each sample and incubated with rotation for 1-3 hours at 4°C. Samples were then washed twice with IP buffer (50 mM Tris–HCl pH 7.4; 150 mM NaCl, 1 mM EDTA) and supernatant was removed. 100 μL 2.5X Laemmli Sample Buffer was then added to the washed beads and denatured at 98°C for 30 minutes.

### Immunoblotting

Samples were run on 4–20% Tris-HCl SDS-PAGE gels (BioRad Criterion) at 90 V for 40 minutes followed by separation at 150 V for 85 minutes. Proteins were transferred to PVDF membranes by electrotransfer (BioRad Criterion Blotter) at 90 V for 2 h. Membranes were blocked in 5% milk in DPBS, 0.1% Tween-20 or 5% BSA in DPBS, 0.1% Tween-20 for 1 h prior to primary antibody incubation overnight at 4°C. Anti-rabbit or anti-mouse IgG horseradish peroxidase (HRP)-conjugated secondary antibodies (1:20000, polyclonal, Jackson ImmunoResearch Laboratories, Cat. Nos. 111-035-003 and 115-035-003) were detected using Pierce™ ECL Western Blotting Substrate (ThermoFisher) on iBright (Thermofisher) blot scanner. Blots were incubated in a 1xPBS, 0.2 M glycine, 1.0% SDS, 1.0% Tween-20, and pH 2.2 stripping buffer before reprobing.

### Primary Antibodies

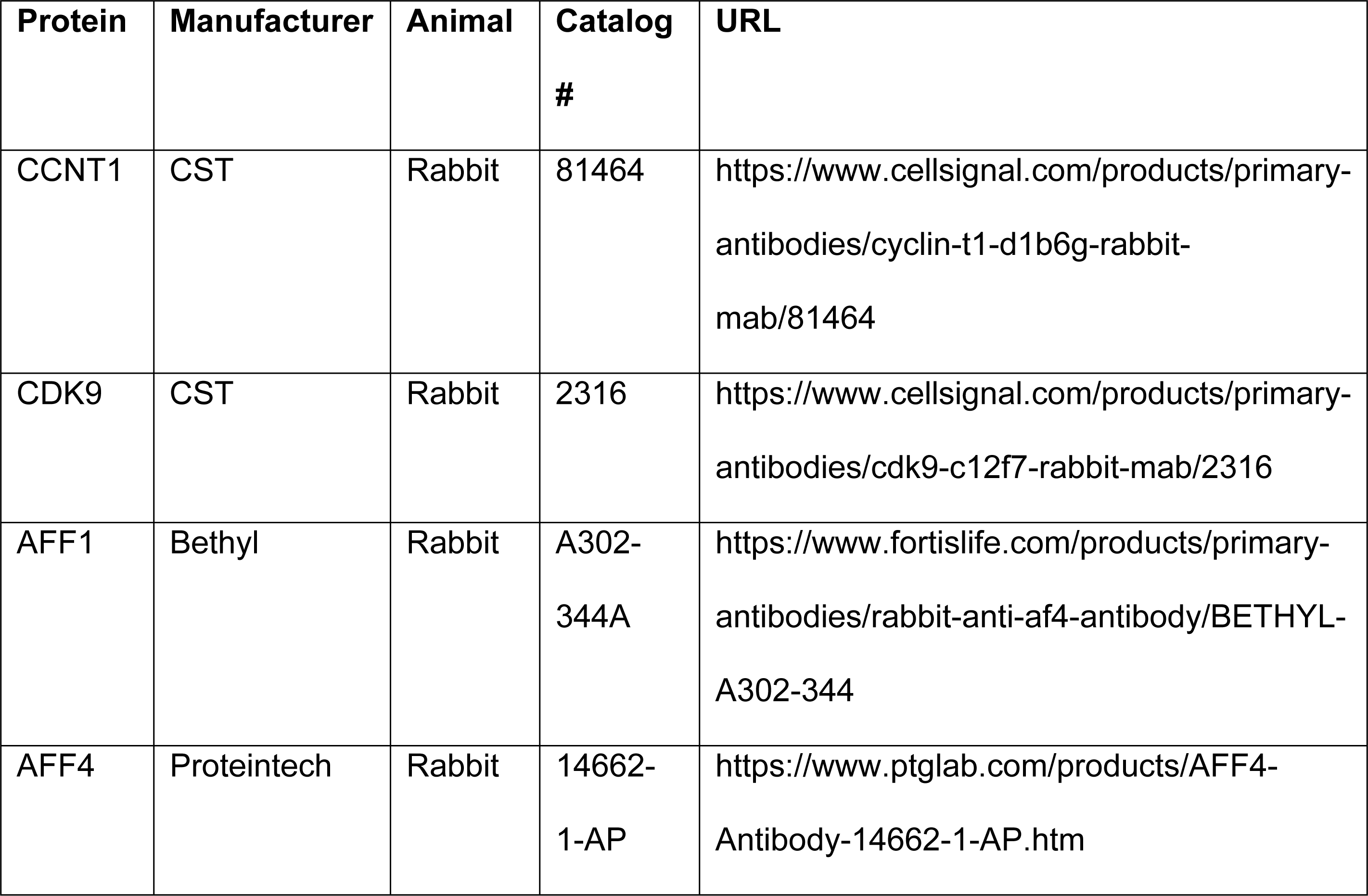
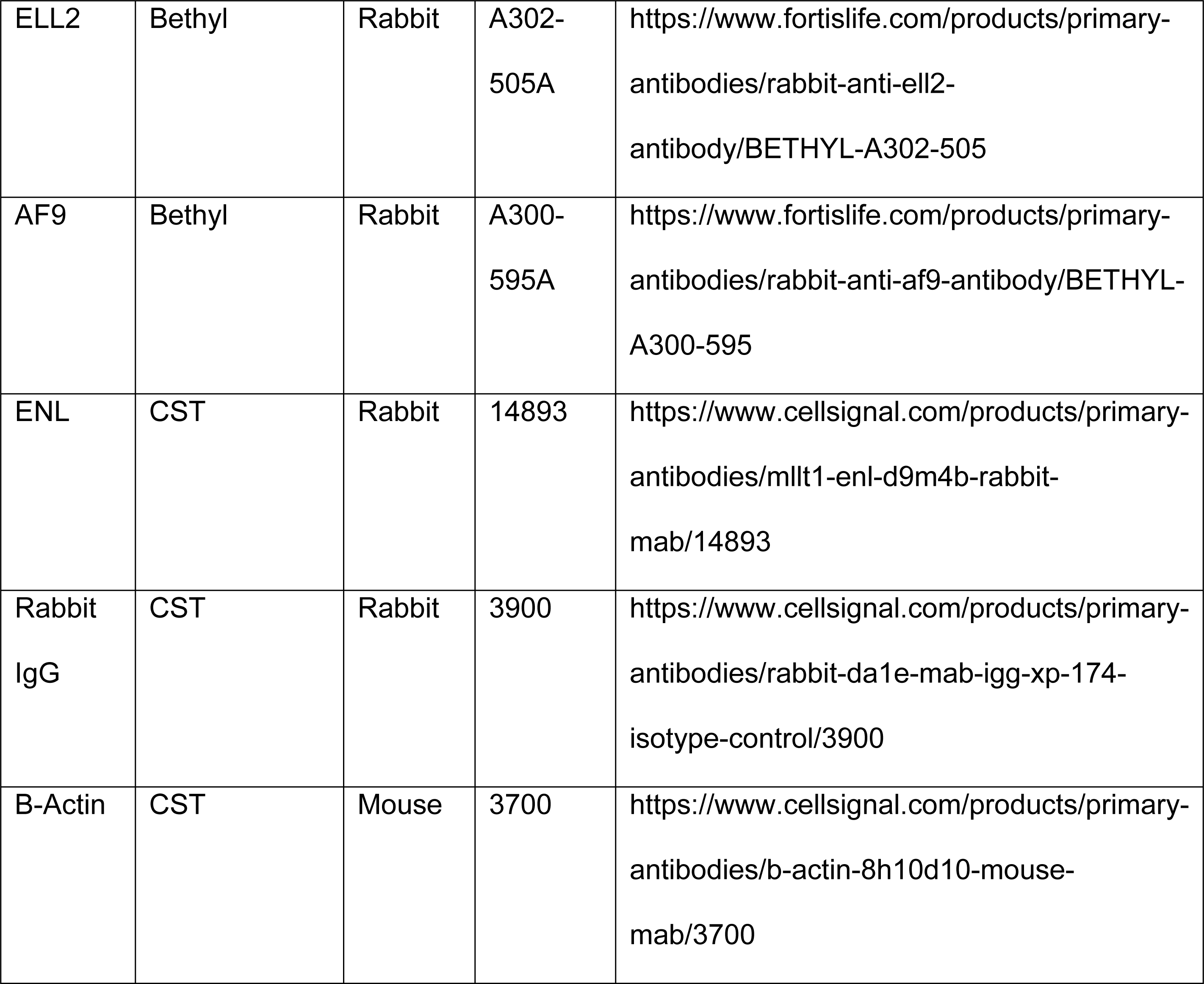

### Preparation of virus stocks for infection of primary CD4+ T cell cultures

Replication-competent reporter virus stocks were generated from an HIV-1 NL4.3 molecular clone in which GFP had been cloned behind an IRES cassette following the viral nef gene (NIH AIDS Reagent Program, catalog no. 11349). Briefly, 10 μg of the molecular clone was transfected (PolyJet; SignaGen) into 5 × 10^6^ human embryonic kidney (HEK) 293T cells (ATCC, CRL-3216) according to the manufacturer’s protocol. Twenty-five milliliters of the supernatant was collected at 48 and 72 hours and then combined. The virus-containing supernatant was filtered through 0.45-mm polyvinylidene difluoride filters (Millipore) and precipitated in 8.5% polyethylene glycol [average molecular weight (Mn), 6000; Sigma-Aldrich] and 0.3 M NaCl for 4 hours at 4°C. Supernatants were centrifuged at 3500 rpm for 20 minutes, and the concentrated virus was resuspended in 0.25 ml of PBS for a 100X effective concentration. Aliquots were stored at −80°C until use.

### HIV-1 Infection of CRISPR-edited primary CD4+ T cell cultures

Edited primary CD4+ T cells were plated into a 96-well, round-bottom plate at a cell density of 1 × 10^5^ cells per well and cultured overnight in 200 μl of complete RPMI 1640 as described above in the presence of IL-2 (20 IU/ml) and 2.5 μl of concentrated virus stock. Cells were cultured in a dark humidified incubator at 37°C and 5% CO_2_. On days 2 and 5 after infection, 75 μl of each culture was removed and mixed 1:1 with freshly made 2% formaldehyde in PBS (Sigma-Aldrich) and stored at 4°C for analysis by flow cytometry. Cultures were supplemented with 75 ml of complete IL-2–containing RPMI 1640 medium and returned to the incubator.

### HIV-1 infection of primary CD4+ T cell cultures treated with KL-2

Activated primary CD4+ T cells were plated into a 96-well, round-bottom plate at a cell density of 1 × 10^5^ cells per well and cultured overnight in 200 μl of complete RPMI 1640 as described above in the presence of IL-2 (20 IU/ml) with different concentrations of KL-2 or equivalent volumes of DMSO. The next day, 2.5 μl of concentrated virus stock was added to each well. Cells were cultured in a dark humidified incubator at 37°C and 5% CO_2_. On days 2 and 5 after infection, 75 μl of each culture was removed and mixed 1:1 with freshly made 2% formaldehyde in PBS (Sigma-Aldrich) and stored at 4°C for analysis by flow cytometry. Cultures were supplemented with 75 ml of complete IL-2–containing RPMI 1640 medium and returned to the incubator.

### Latent cell line drug treatment assays

J-Lat 5A8, J-Lat 11.1, J-Lat 6.3, J-Lat A72, U1, and ACH-2 cell lines were plated in 96-well flat bottom plates at a density of 50,000 cells/200 μl supplemented RPMI 1640. Cells were DMSO-treated or treated with KL-2 (6.25 μM), JQ1, PMA, and AZD5582 for 48 hours. Cells were then washed in PBS and resuspended in PBS + 1% formaldehyde and fixed for 30 minutes. Analysis was performed by flow cytometry gating on GFP-positive cells for J-Lat cell lines.

| Cell Line | JQ1 | PMA | AZD5582 |
| --- | --- | --- | --- |
| J-Lat 5a8 | 1 μM | 0.5 nM | 10 nM |
| J-Lat 6.3 | 1 μM | 0.5 nM | 10 nM |
| J-Lat 11.1 | 1 μM | 0.5 nM | 10 nM |
| J-Lat A72 | 1 μM | 0.5 nM | 10 nM |
| U1 | 1 μM | 0.5 nM | 500 nM |
| ACH-2 | 1 μM | 1 nM | 500 nM |

### Immunostaining

Extracellular staining was performed on uninfected, live primary T cell populations with anti-CD4-PE (Miltenyi Biotec, 130-113-225), anti-CXCR4-APC (Miltenyi Biotec, Cat#130-120-708), and anti-CD25-APC (Miltenyi Biotec, Cat#1130-115-535) antibodies according to the manufacturer’s instructions. Briefly, cells were pelleted, and media was removed. Cells were then washed once with DPBS and resuspended in a 1:50 dilution of the appropriate antibodies in MACS buffer (PBS + 0.5% bovine serum albumin (BSA) and 2 mM EDTA) and incubated for 15 minutes at 4°C. Cells were then pelleted, washed with MACS buffer, and suspended in PBS+ 1% formaldehyde for 30 minutes prior to flow cytometry.

Viability staining was performed on various cell populations using the amine-reactive dye Ghost Red 710 (Tonbo, Cat#13-0871-T100) according to manufacturer’s instructions. Briefly, cells were pelleted and media was removed. Cells were then washed once with DPBS and resuspended in a ghost red dye solution consisting of 1 μL of ghost red dye per 1 mL of DPBS and incubated for 30 minutes at 4°C. Cells were then pelleted, washed with MACS buffer, and suspended in DPBS + 1% formaldehyde for 30 minutes prior to flow cytometry.

Intracellular HIV-1 p24 staining was performed on fixed U1 and ACH-2 cells with KC57-FITC (Beckman Coulter, Cat#6604665). After allowing at least 30 minutes for fixation, cells were pelleted and washed with DPBS. Subsequently, cells were suspended in DPBS plus 1% BSA, 0.1% w/v saponin and incubated at room temperature for 20 minutes to block and permeabilize. Cells were then spun down to remove the supernatant and incubated for 30 minutes at room temperature in the dark with KC57-FITC at a concentration of 1:50 in DPBS plus 1% BSA, 0.1% saponin w/v. Cells were then pelleted again, washed with PBS plus 1% BSA, and suspended in 1% formaldehyde PBS for fixation prior to flow cytometry.

### Flow cytometry and analysis of viability/infection/reactivation data

Flow cytometry analysis was performed on an Attune NxT acoustic focusing cytometer (Thermo Fisher Scientific), recording all events in a 40-μl sample volume after one 150 μl of mixing cycle. Data were exported as FCS3.0 files using Attune NxT Software v3.2.0 and analyzed with a consistent template on FlowJo. Briefly, cells were gated for lymphocytes by light scatter followed by doublet discrimination in both side and forward scatter. Cells with equal fluorescence in the BL-1 (GFP) channel and the VL-2 (AmCyan) channel were identified as autofluorescent and excluded from the analysis. A consistent gate was then used to quantify the fraction of remaining cells that expressed the target of interest.

### RNA Isolation and cDNA synthesis

Total RNA isolation from cells (typically 100,000 cells) treated with our LRA panel in the presence or absence of KL-2 was carried out with an RNeasy kit (Qiagen), with the optional on-column deoxyribonuclease I digestion step. The isolated total RNA was eluted in ribonuclease-free water and RNA concentrations were subsequently quantified using a NanoDrop One (Thermo Fisher). cDNA was synthesized from extracted RNA (typically 100 ng) using the SuperScript IV first strand synthesis system (Invitrogen, Cat#18091050) according to the manufacturer’s instructions. Briefly, a 13 μl reaction mixture was made with random hexamers (1 μl; 50 ng/ μl), 10 mM dNTP (1 μl), template RNA (100 ng) and DEPC-treated water. The RNA-hexamer mix was then incubated at 65°C for 5 minutes before incubating on ice for another minute. Then a mix of 5x SSIV Buffer (4 μl), 100 mM DTT (1 μl), Ribonuclease Inhibitor (μl), and SuperScript IV Reverse Transcriptase (1 μl, 200 u/ μl) was added to each sample. The combined reaction mixture was then incubated at 23C for 10 minutes, followed by 50°C for 10 minutes, and the reaction was inactivated by incubation at 80C for 10 minutes. Following complete reverse transcription, residual RNA was cleared by the addition of 1 μl *E. Coli* RNase H to each reaction mix and incubated at 37°C for 20 minutes. The RT products were then stored at -20°C.

### Analysis of HIV-specific RNA transcripts

Following cDNA synthesis, viral transcripts were assessed by qRT-PCR as described previously (13). For HIV-1 TAR, the following primers were used: PF: 5′-GTCTCTCTGGTTAGACCAG-3′; PR: 5′-TGGGTTCCCTAGYTAGCC-3′; and probe: 5′-AGCCTGGGAGCTC-3′. HIV-1 Long LTR levels were assessed using the following primers: PF: 5′-GCCTCAATAAAGCTTGCCTTGA-3′; PR: 5′-GGGCGCCACTGCTAGAGA-3′; and probe: 5′-CCAGAGTCACACAACAGACGGGCACA-3. For expression level normalization, *β-Actin* was used (Thermo Fisher Scientific, catalog no. 4331182). The reaction was performed using TaqMan Fast Advanced Master Mix (Thermo Fisher Scientific, catalog no. 4444553) according to manufacturer’s instructions. Briefly, a 10 μl of reaction was mixed using 5 μl of Taqman Master Mix, 0.5 μl of 20x primer probe mix (18 μM of primers and 5 μM probe), 2.5 μl water, and 2 μl of template cDNA. The PCR cycles were as follows: 50°C for 2 minutes, 95°C for 20 seconds, followed by 40 cycles of 95°C for 1 second and 60°C for 20 seconds.

### Tat-Rescue of U1 cells

Codon optimized HIV-1 NL4.3 Tat with a C-terminal 2xStrep(TagII)-TEV-3xFlag tag was ordered as a GeneBlock (IDT) and inserted into BamHI/ECORI linearized pLVX-TetOne-Puro by Gibson Assembly and subsequently sequence verified. Briefly, 10 μg of the molecular clone was transfected (PolyJet; SignaGen) into 5 × 10^6^ human embryonic kidney (HEK) 293T cells (ATCC, CRL-3216) according to the manufacturer’s protocol. Twenty-five mL of the supernatant was collected at 48 and 72 hours and then combined. The virus-containing supernatant was filtered through 0.45-mm polyvinylidene difluoride filters (Millipore) and precipitated in 8.5% polyethylene glycol [average molecular weight (Mn), 6000; Sigma-Aldrich] and 0.3 M NaCl for 4 hours at 4°C. Supernatants were centrifuged at 3500 rpm for 20 minutes, and the virus was resuspended in 0.25 ml of PBS for a 100X effective concentration. Aliquots were stored at −80°C until use. Aliquots were thawed and added to cultures of U1 cells at a concentration of 1:100 in complete RPMI and cells were cultured for 48 hours. Following transduction, the cells were pelleted, and media was removed and replaced with fresh complete RPMI + 10 μg/mL of puromycin to select for successfully transduced cells. Cells were cultured in selective growth media for 7 days. After selection, cells were plated in 96-well flat-bottom plates according to previous methods and treated with latency reversing agents and doxycycline for 48 hours before analysis of p24 expressing cells by flow cytometry.

### Treatment of patient PBMCs

*Study subjects*: We selected five study subjects enrolled in the Northwestern University Clinical Research Site for the MWCCS who were well-suppressed and had received antiretroviral drugs for at least 5 years. We obtained patient PBMCs from cryostorage with characteristics more than 5 years after starting antiretroviral therapy and undetectable plasma HIV-1 (<50 copies/ml). Laboratory procedures for clinical sample management are described previously (65). The Institutional Review Board of Northwestern University approved the study (STU00022906-CR0008) with most recent approval date of 16 May 2022. All participants provided written informed consent.

*Total RNA isolation*: Total RNA was isolated from cultures of 1 x 10^6^ PBMCs that were treated with for 48 hours with JQ1, or AZD5582 in the presence or absence of KL-2 using an RNeasy kit (Qiagen), with the optional on-column deoxyribonuclease I digestion step. The isolated total RNA was eluted in ribonuclease-free water, cleaned, and concentrated using the RNA Clean and Concentration kit (Zymo) and assessed for quantity and quality by Qubit (Thermo Fisher Scientific) and 4200 TapeStation (Agilent), respectively before qRT-PCR.

*Validation of cell-associated HIV-1 RNA*: We performed real-time qRT-PCR using an HIV-1-gag– specific primers-probe (FAM) set: HIV-1-gagF, 5′-GGTGCGAGAGCGTCAGTATTAAG-3′; HIV-1-gagR, 5′-AGCTCCCTGCTTGCCCATA-3′; HIV-1-gagProbe, 6FAM-5′-TGGGAAAAAATTCGGTTAAGGCCAGGG-3′-QSY. We used the lactate dehydrogenase A (LDHA) gene for the internal normalization primers-probe (VIC) set (VIC-MGB: assay ID Hs03405707_g1; TaqMan Gene Expression Assay, Thermo Fisher Scientific). Briefly, a 10-μl RT-PCR mixture contained TaqMan Fast Virus 1-Step Master Mix, 400 nM forward and reverse HIV-1-gag primers, 0.3 μl of LDHA Gene Expression Assay (Thermo Fisher Scientific), 250 nM each of the probes, and 5 μl of extracted RNA or water for the no template controls. We programmed the 7900HT real-time PCR system (Applied Biosystems) for 20 minutes at 50°C and 20 seconds at 95°C, followed by 40 cycles of 15 seconds at 95°C and 60 seconds at 60°C. The qRT-PCR data were analyzed in technical triplicate. We calculated the fold change in gene expression using the standard 2-ΔΔCT method.

### Statistical Analysis

All statistical analysis was performed using GraphPad Prism version 10.2.0 (392) for Windows 64-bit, GraphPad Software, Boston, Massachusetts USA, www.graphpad.com”

## ACKNOWLEDGEMENTS

We would like to thank members of the Hultquist Lab for comments, manuscript revision, and helpful discussions. We would like to thank Cristina Vaca and the laboratory of Dr. Richard T. D’Aquila (Northwestern University) for sharing the J-Lat 6.3 and J-Lat 11.1 cell lines. Additionally, we would like to thank Dr. Elena Martinelli (Northwestern University) for sharing the U1 cell line. This research was supported by NIH/NIAID funding for the HIV Accessory & Regulatory Complexes (HARC) Center (U54 AI170792, J.F.H.), Northwestern University Cell and Molecular Basis of Disease (CMBD) Training Program (T32 GM008061, W.J.C.), NIH funding for the Third Coast Center for AIDS Research (P30 AI117943, J.F.H.), and NIH/NIAID grants for HIV research (R01AI176599, R01AI167778, R01AI150455, R01AI165236, R01AI150998, R21AI174864, and R56AI174877 to J.F.H.).

## AUTHOR CONTRIBUTIONS

Conceptualization: W.J.C., A.S., J.F.H.; Methodology: W.J.C., L.M.S., J.F.H.; Validation: W.J.C.; Formal Analysis: W.J.C., E.-Y.K.; Investigation: W.J.C., L.M.S., D.C., M.W., A.W.H., S.H.A.S.; Resources: L.M.S., S.H.A.S., A.S., J.F.H.; Data Curation: W.J.C.; Writing – Original Draft: W.J.C., J.F.H.; Writing – Review and Editing: W.J.C., E.-Y.K., S.M.W., J.F.H.; Visualization: W.J.C.; Supervision: E.-Y.K., S.M.W., A.S., J.F.H.; Project Administration: L.M.S., E.-Y.K., S.M.W., A.S., J.F.H.; Funding Acquisition: S.M.W., A.S., J.F.H.

## SUPPLEMENTAL FIGURES AND LEGENDS

**Supplemental Figure 1.**
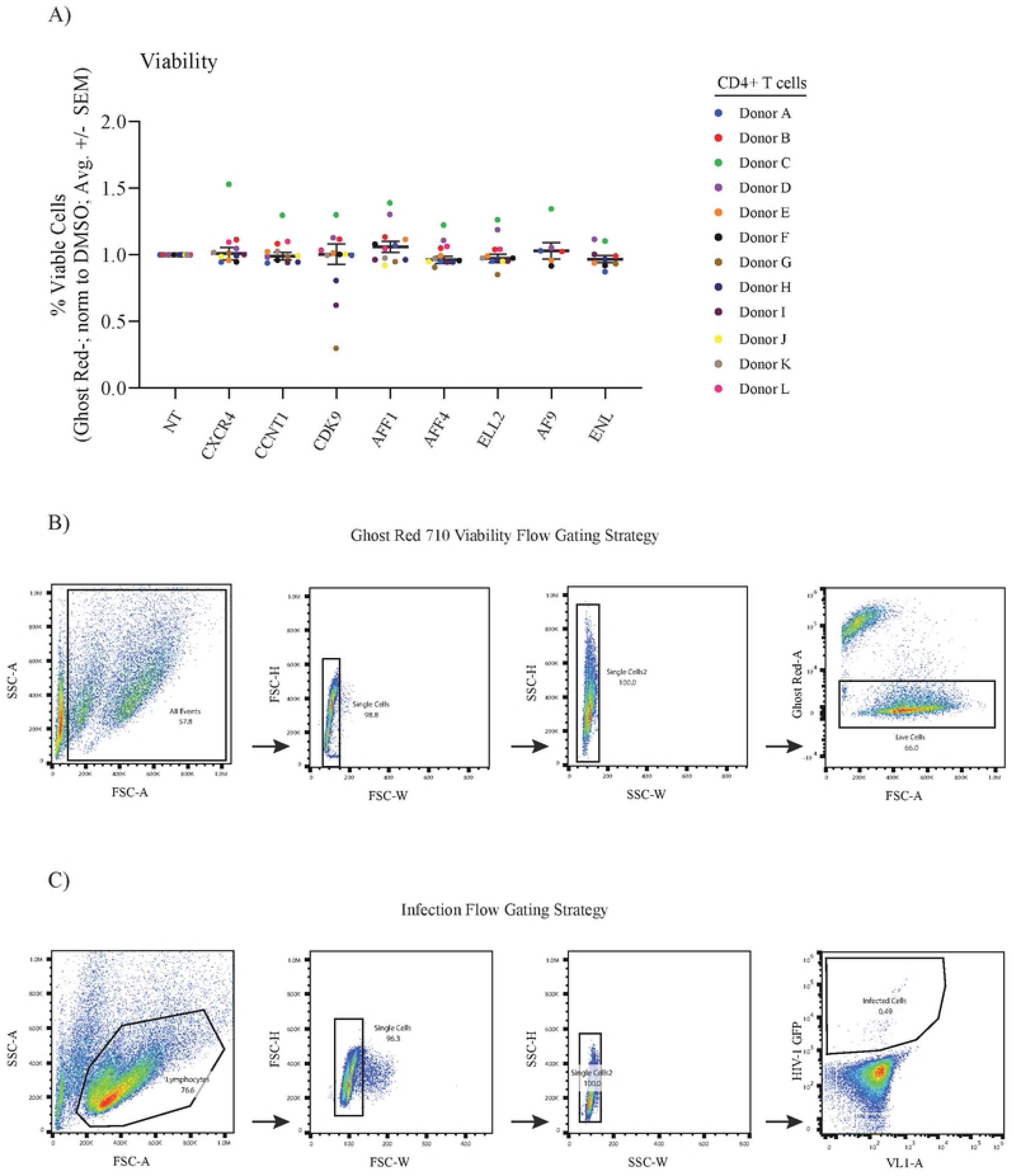
Primary cell viability and gating strategy. **A)** Percent viable primary CD4+ T cells (normalized to the donor-matched NT control) 72 hours after electroporation with multiplexed CRISPR-Cas9 RNPs targeting the indicated genes as measured by amine dye staining and flow cytometry. Each dot represents the average of technical triplicates; the black line represents the mean of means ± standard error. n = 12 donors for NT, CXCR4, CCNT1, CDK9, AFF1, AFF4, and ELL2; n=6 donors for AF9; n=9 donors for ENL. Statistics were calculated by 2-way ANOVA with Dunnet’s Multiple Comparison Test; no significant differences were observed. **B)** Gating strategy for quantification of percent viable primary CD4+ T cells via sequential application of a live cell gate, two single-cell gates, and a fluorophore gate (FlowJo v10.7.1). **C)** Gating strategy for quantification of percent HIV-1 infection in primary CD4+ T cells via sequential application of a live cell gate, two single-cell gates, and autofluorescence exclusion (FlowJo v10.7.1).

**Supplemental Figure 2.**
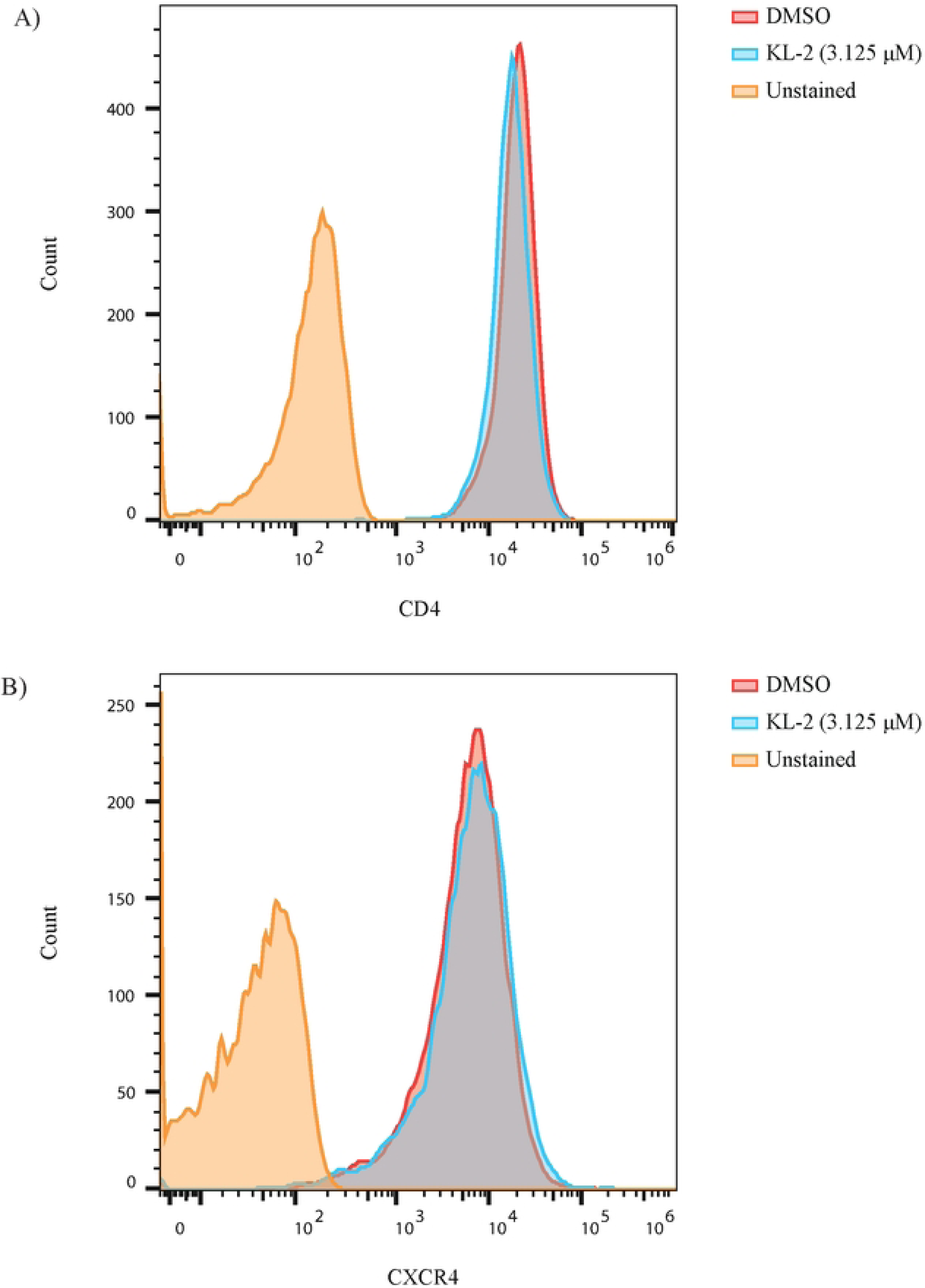
Histograms of CD4 and CXCR4 expression on primary CD4+ T cells after KL-2 treatment. **A)** Histogram of cell surface CD4 expression on activated CD4+ T cells treated with DMSO or 3.125 μM KL-2 for 48 hours as measured by immunostaining and flow cytometry (one representative donor, visualized in FlowJo v10.7.1). **B)** Histogram of cell surface CXCR4 expression on activated CD4+ T cells treated with DMSO or 3.125 μM KL-2 for 48 hours as measured by immunostaining and flow cytometry (one representative donor, visualized in FlowJo v10.7.1).

**Supplemental Figure 3.**
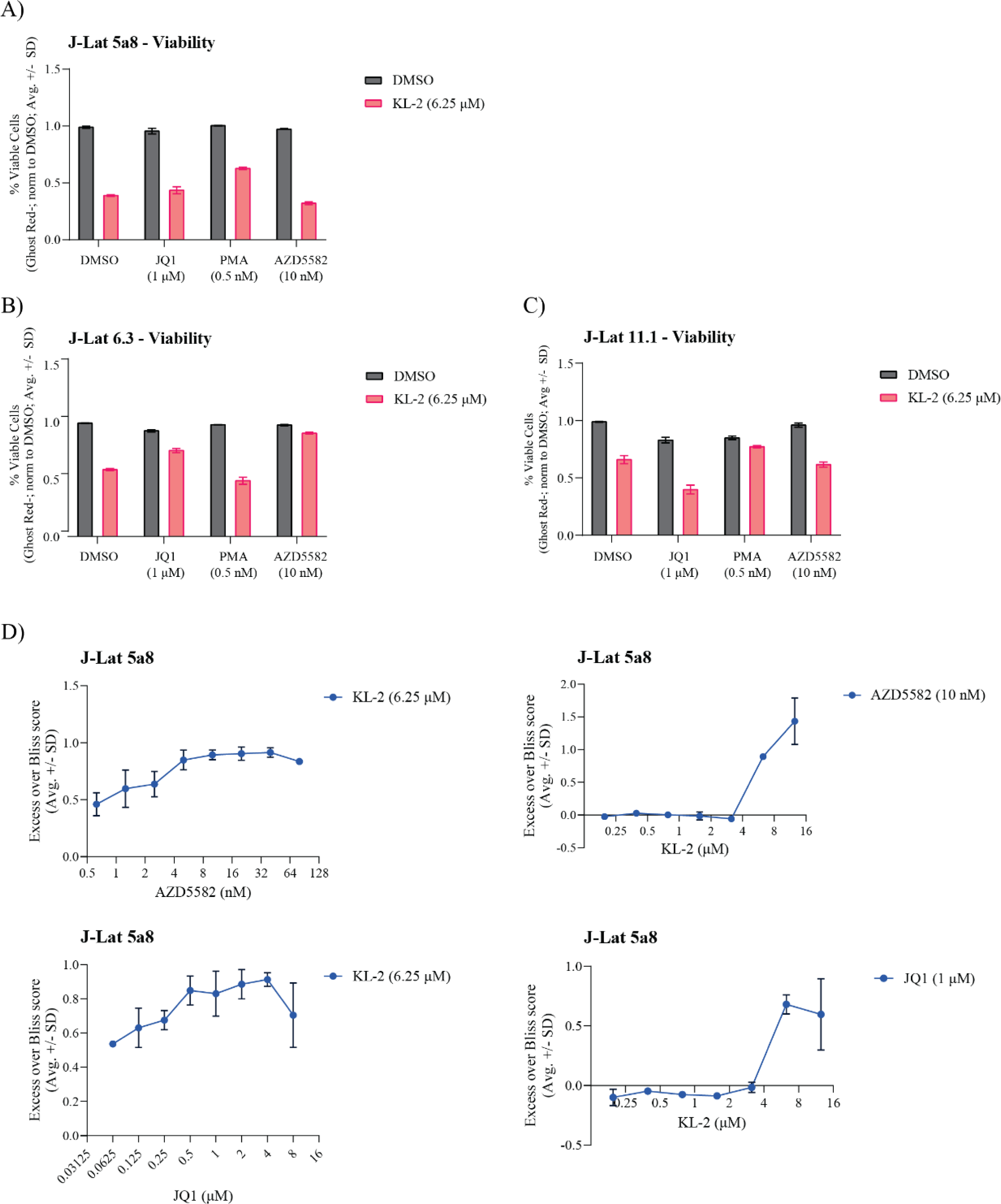
J-Lat cell viability upon LRA treatment and reactivation. **A)** Percent viable J-Lat 5A8 cells (normalized to the DMSO treated control) 48 hours after treatment with the indicated LRAs in the presence and absence of 6.25 μM KL-2 as measured by amine dye staining and flow cytometry. Each bar represents the average ± standard deviation of technical triplicates. The same data are shown for **B)** J-Lat 6.3 cells and **C)** J-Lat 11.1 cells. **D)** Excess over Bliss scores for latency reactivation in J-Lat 5a8 cells treated with combinations of KL-2 and AZD5582 (top) or JQ1 (bottom). Cells were treated in biological duplicates with the mean excess over Bliss score +/- SD between replicates shown.

**Supplemental Figure 4.**
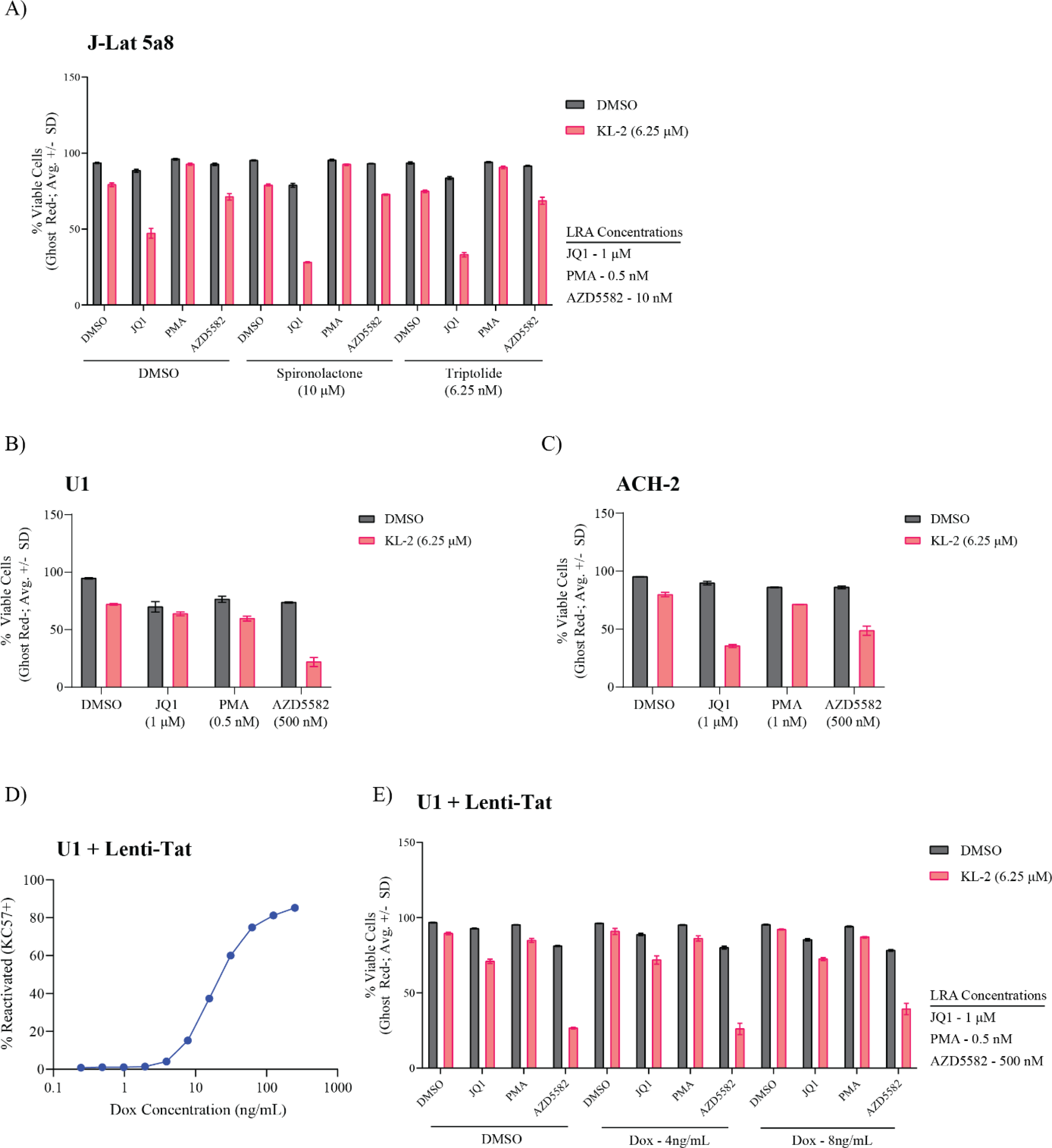
J-Lat, U1, and ACH-2 cell viability upon LRA treatment and reactivation. **A)** Percent viable J-Lat 5A8 cells 48 hours after treatment with the indicated LRAs in the presence and absence of 6.25 μM KL-2 and the Tat inhibitors Spironolactone and Triptolide as measured by amine dye staining and flow cytometry. Each bar represents the average ± standard deviation of technical triplicates. **B)** Percent viable U1 cells 48 hours after treatment with the indicated LRAs in the presence and absence of 6.25 μM KL-2 as measured by amine dye staining and flow cytometry. Each bar represents the average ± standard deviation of technical triplicates. **C)** Percent viable ACH-2 cells 48 hours after treatment with the indicated LRAs in the presence and absence of 6.25 μM KL-2 as measured by amine dye staining and flow cytometry. Each bar represents the average ± standard deviation of technical triplicates. **D)** Percent reactivated (KC57-FITC+) Lenti-Tat U1 cells after 48 hours of treatment with increasing concentrations of doxycycline. **E)** Percent viable Lenti-Tat U1 cells 48 hours after treatment with the indicated LRAs in the presence and absence of 6.25 μM KL-2 and differing amounts of doxycycline as measured by amine dye staining and flow cytometry. Each bar represents the average ± standard deviation of technical triplicates.

